# UG/Abi: a highly diverse family of prokaryotic reverse transcriptases associated with defense functions

**DOI:** 10.1101/2021.12.02.470933

**Authors:** Mario Rodríguez Mestre, Linyi Gao, Shiraz A. Shah, Adrián López-Beltrán, Alejandro González-Delgado, Francisco Martínez-Abarca, Jaime Iranzo, Modesto Redrejo-Rodríguez, Feng Zhang, Nicolás Toro

## Abstract

Reverse transcriptases (RTs) are enzymes capable of synthesizing DNA using RNA as a template. Within the last few years, a burst of research has led to the discovery of novel prokaryotic RTs with diverse antiviral properties, such as DRTs (Defense-associated RTs), which belong to the so-called group of unknown RTs (UG) and are closely related to the Abortive Infection system (Abi) RTs. In this work, we performed a systematic analysis of UG and Abi RTs, increasing the number of UG/Abi members up to 42 highly diverse groups, most of which are predicted to be functionally associated with other gene(s) or domain(s). Based on this information, we classified these systems into three major classes. In addition, we reveal that most of these groups are associated with defense functions and/or mobile genetic elements, and demonstrate the antiphage role of four novel groups. Besides, we highlight the presence of one of these systems in novel families of human gut viruses infecting members of the Bacteroidetes and Firmicutes phyla. This work lays the foundation for a comprehensive and unified understanding of these highly diverse RTs with enormous biotechnological potential.

## Introduction

Reverse transcriptases (RTs, also known as RNA-directed DNA Polymerases) are enzymes present in all three domains of life whose main function is to polymerize DNA strands using RNA as a template. Although they were first discovered by Temin & Baltimore in 1970 (1,2), prokaryotic RTs were not observed until 1989 when they were found to be the main component of retrons (3). Later research revealed that most reverse transcriptases (80%) can be phylogenetically clustered into three major lineages: group II introns, diversity-generating retroelements (DGRs), and retrons, which are the best known due to their ecological implications and biotechnological applications. Other minor clades of RTs include abortive infection (Abi) RTs, CRISPR-Cas-associated RTs, Group II-like (G2L), the unknown groups (UG), and *rvt* elements (4–6).

Comprehensive and systematic analysis of prokaryotic RTs (7), identified the association of RTs with CRISPR-Cas systems and 5 novel gene families (D, E, F1, F2, and G, now known as UG9, UG6, UG1, UG5, and UG3+UG8, respectively). Further research revealed the existence of other uncharacterized RTs from distinct clades (UG1-UG16 and Group II-like, including those described by Kojima & Kanehisa) (6,8,9), and a more recent work disclosed 11 additional UG RT groups (UG17-UG28), pointing out that UG RTs and Abi RTs may form a novel major lineage branching off from a common node (5). Although it was initially thought that UG and Abi RTs were not very common, it is known that they represent at least 11% of all prokaryotic RTs, showing an enormous diversity and holding great promise for the development of new biotechnological tools (4).

Abi systems can function as prokaryotic defense mechanisms against certain phages (10,11). They are generally constituted by a sensing module that recognizes a phage-specific signal and an effector module that generates a response, either by blocking the viral infection cycle, halting host metabolism, or causing cell death (12). Although there are more than 20 different Abi systems, only a few have been well characterized. Among them, AbiA, AbiK, and AbiP2 share an N-terminal RT domain (13–17), with AbiA harboring an additional C-terminal HEPN domain (higher eukaryotes and prokaryotes nucleotide-binding domain) with predicted RNAse activity (18). Both AbiA and AbiK are commonly found in plasmids and have been shown to protect *Lactococcus* spp. from diverse phage infections (13), whereas AbiP2 is commonly found in a hypervariable region of P2 prophages in *E. coli* and confers resistance against T5 phage (17). It has been hypothesized that AbiA and AbiK could have a similar mechanism of action, as both of them confer protection against the same phages either by blocking DNA replication or targeting functionally-related proteins (11). Furthermore, phages escaping AbiK and AbiA interference have been shown to harbor point mutations in single-strand annealing proteins (SSAPs) involved in DNA replication (19). AbiK is the best characterized, and it has been hypothesized to have protein-primed untemplated DNA-polymerase activity (14). The residues responsible for this activity are thought to be located at the C-terminal region (14) that along with the RT domain, is essential for its biological role (13,20).

Although Abi and UG RTs appear to be phylogenetically related, they were thought to be functionally unrelated as they bear distantly related RT domains. Also, previous analyses pointed out the high divergence at the sequence level between AbiK, AbiA, and AbiP2 (6). That notwithstanding, recent research that employed a systematic methodology to search for novel antiphage systems (21) highlighted that some members of UG RTs (UG1, UG2, UG3-UG8, UG15, UG16, named DRTs type 1-5 respectively for Defense-associated RTs) act as defense mechanisms against bacterial viruses. This suggests a functional link between Abi RTs and UG RTs and supports the idea that different families of RTs may be implicated in immunity against bacteriophages (4).

Even though some UG/Abi RTs members have been reported to have antiphage functions and associated domain(s) required for this function, some others remain poorly characterized due to insufficient information on their genomic context, associated genes, or biological roles. Considering the great diversity of UG/Abi RTs, their possible common origin, and the recently disclosed role of retrons and DRTs (21–24), we hypothesize that the UG/Abi RT lineage may constitute a novel family of defense-related RTs with high divergence and a plethora of associated genes. In this work, we performed a systematic analysis of UG/Abi RTs and their neighborhood in search of associated genes and defense hallmarks. As a result, we expanded the number and diversity of UG/Abi RTs with novel groups, of which most are associated with other protein domain(s) and located within defense islands/hotspots. Based on this information, the UG/Abi RTs could be classified into three major classes, a first class of RTs fused to HEAT-like repeats, a second class of highly diverse RTs not fused to any known domain, and a third class commonly associated with C-N hydrolase (carbon-nitrogen hydrolase, also known as nitrilase) or phosphohydrolase domains. Besides, we demonstrate the antiphage activity of three Class 1 members and an additional Class 2 member. Moreover, we reveal that UG27, a Class 2 UG/Abi RT, is commonly encoded in several groups of predicted human gut viruses infecting members of the *Bacteroidetes* and *Firmicutes* phyla, which encode a putative non-coding RNA (ncRNA) with a common secondary structure. Finally, different UG groups (UG2, UG3+UG8 and UG28) that have been described to possess antiphage properties (DRT type 2, DRT type 3 and DRT type 9 respectively) (21) also encode ncRNAs with conserved secondary structures, thought to be essential for the functioning of the systems. Altogether, these findings reveal that the UG/Abi RTs family is a highly diverse and widespread lineage of prokaryotic reverse transcriptases associated with defense functions that would play a very important role in virus-host conflicts.

## Methods

### Construction of a comprehensive dataset of representative UG/Abi RTs dataset

To increase the number of UG/Abi RT sequences, the most up-to-date phylogenetic tree of prokaryotic RTs based on an alignment of the RT domain of 9141 RTs (5) was used as a reference. Custom HMM profiles for every phylogenetic group (group II introns, retrons, DGRs, CRISPR-associated RTs, G2L, and Abi/UG RTs) were built using hmmbuild from the suite HMMER 3.3 (25). Then, the NR database (ftp://ftp.ncbi.nlm.nih.gov/blast/db/FASTA/) was searched (February 2021) with all profiles using hmmsearch (E-value <1e-05) (25). Only sequences in which the E-value using the UG/Abi profile was at least one order of magnitude lower than the E-value using other profiles were kept. After this, sequences present in Eukaryotic genomes or with unassigned taxonomy were removed using the NCBI API (26), which resulted in a dataset containing 12, 209 potential UG RT sequences present in Bacteria, Archaea, and their viruses. To remove sequence redundancy, the 12209 sequences were clustered using CD-HIT (27) with default parameters and the option -c 0.85 (85% AAI; average amino acid identity), which resulted in 5727 clusters that were further filtered based on quality and contig completeness criteria to perform a neighborhood analysis. Sequences with no associated NCBI nucleotide accession were discarded from the analysis. Further, the NCBI Identical Protein Groups (IPG) database (26) was used to retrieve information about the completeness of the assemblies in which each of the sequences was found. For each entry, the best IPG candidate was selected based on reference assignments and/or completeness (reference genomes > complete genomes > scaffolds/contigs). If not found in any complete genome, sequences in contigs with the greater neighborhood information (i.e., large contigs where the RT is not located close to their ends) were prioritized by choosing those with the highest W value, a parameter that illustrates the length of the contig weighted by how centered the ORF is in the contig; described below.

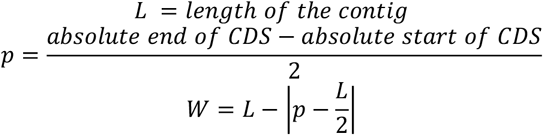

### Phylogenetic and network analysis of UG/Abi RTs

The cd-hit-2d tool (27) was used to label sequences highly similar (>95% AAI) to the reference entries. After this, a multiple sequence alignment (MSA) of RT1-7 motifs was performed using the MAFFT software (28) with default parameters, and a phylogenetic tree was built using FastTree (29) with the WAG evolutionary model, and discrete gamma model with 20 rate categories. A phylogenetic tree was also constructed with IQ-TREE v1.6.12, with 1000 ultra-fast bootstraps (UFBoot) and SH-like approximate likelihood ratio test (SH-aLRT) with 1000 replicates (-bb 1000 -alrt 1000 options) (30), using the LG+R10 model identified as the best model by Modelfinder (31). To compare full-length sequences, a sequence-similarity network (SSN) of these sequences was also built using the EFI-EST resource (32) and visualized using Cytoscape (33) with the force-directed layout using the BLAST score as a weight.

### Retrieval and clustering of neighbor proteins

Coding sequences (CDS) located within ± 10 kb of the start and the end of our query proteins were retrieved using the feature table resource from the NCBI Entrez API (26). Due to the frequent misannotation of ORFs, nucleotide sequences of the intergenic regions were also retrieved, and ORFs were predicted using the Prodigal tool with -c -m -n -p meta parameters (34). Neighbor CDS and ORFs predicted in the intergenic sequences were joined and clustered using MMseqs2 (35) with a profile-based deep clustering method previously described (36) using 10 iterations, which rendered 3240 neighbor clusters.

### Prediction and annotation of functionally associated genes

Genes functionally associated with UG/Abi RTs were predicted using a methodology previously described (36). Briefly, a presence/absence matrix of neighbor clusters surrounding UG/Abi RT entries was analyzed in search of non-random patterns of association. Based on the distribution of clusters across the tree and the average amino acid identity (AAI) of the co-located RTs, 193 out of 3240 protein clusters with more than 5 members were selected as potentially linked. After this, MSAs and HMM profiles were built using MAFFT (28) with default parameters and hmmbuild (25), respectively. Domain annotation of protein clusters was done using HHsearch against the hh-formatted PFAM (37), CDD (38), COG (39), ECOD (40), and pdb30 databases jointly distributed with HH-suite (41) (Supplementary Table S2). In addition, we also performed comparisons against profiles built using the eggNOG (Bacteria, Archaea and Viruses) (42), pVOG (43), and mMGE (44) databases.

### Group adscription and refinement of UG/Abi RTs

Group adscription of sequences was manually assigned based on the phylogeny, the sequence-similarity network, the presence of labeled reference sequences, and the presence/absence matrix. Individual sequences that were difficult to classify, with low support, or with little information about the neighborhood were manually removed. After performing this task iteratively, and rebuilding MSAs, phylogenetic trees, and SSNs as described above, 5022 UG/Abi RT bona fide representative sequences were retained. Those groups that did not include previously described or unmatched sequences were treated as novel groups.

### Sequence and structure-based annotation of UG/Abi RTs domains

For every UG/Abi RT group, MSAs were built using MAFFT-einsi (28) with default parameters. Groups with bimodal length distribution were subdivided into small and large variants, and MSAs were built independently. Annotation of UG/Abi RTs was done using hhsearch and PFAM, COG, CDD, and ECOD databases. We then performed structural predictions employing trRosetta (45) using previously built MSAs as input for modeling. After this, predicted models were compared against the PDB database using the DALI webserver (46). For αRep domains found in Class 1 UG/Abi RTs, motif boundaries were obtained from trRosetta using contact maps and predicted structure models as a reference. Then, trimmed MSAs were used as a query to perform further trRosetta structure predictions. A multiple protein structure alignment was built for every Class1 predicted repeated structure model, by using the mTM-align web server (47) and PyMOL Molecular Graphics System (Schrödinger, LLC) Cealign command with UG8 αRep domain as an anchor.

### Taxonomy assignment

To determine the taxonomic distribution of every UG/Abi RT group, every representative sequence was queried against the NCBI taxonomy database (26), and information about the domain, phylum, class, order, family, and genus was retrieved for every associated genome (Supplementary Table S1). Then, relative abundances of phyla across the different UG/Abi RT groups were calculated and plotted using the ggplot2 R package (48). For every group, those phyla with <1% of relative abundance were removed to improve the visualization.

### Prediction of RT defense association

Defense genes were predicted as previously described (21). Briefly, a total of 174, 080 bacterial and archaeal genomes were downloaded from GenBank in 2018, and highly similar proteins (at least 98% sequence identity and coverage) were discarded using the linclust option in MMseqs2 with parameters --min-seq-id 0.98 -c 0.98. To identify homologs of each of the 42 groups of RTs within this data set, representatives from each RT group (5, 022 total sequences) were used as search seeds for MMseqs2. To be labeled as an RT homolog, proteins were required to have a minimum of 80% sequence identity and 80% coverage (--min-seq-id 0.8 -c 0.8) to at least one RT search seed. Following homolog identification, the defense association frequency (defined as the proportion of homologs within 5 kb or 5 ORFs from the nearest annotated known defense system) for each RT group was calculated as previously described (21) (Supplementary Table S3). Some groups had inconclusive defense association frequencies or a low number of homologs (less than 50). Because of this, they were manually examined for signatures of strong defense association (e.g. operonization with a type I restriction-modification system) as further evidence in support of a defense prediction (Supplementary Figure S2).

### Cloning

Genes were chemically synthesized as gene fragments (GENEWIZ) or amplified with Q5 (New England Biolabs) or Phusion Flash (Thermo Scientific) polymerase. The native promoter and ORF sequences were retained in all cases except for UG15, which was recoded (21). Inserts were cloned into a low copy pACYC184-derived empty vector (Addgene # 157879) between the HindIII and EcoRI restriction sites using NEBuilder HiFi DNA Assembly mix (New England Biolabs). The full sequences of all plasmids were verified as previously described (21,49). Briefly, ∼25ng of each plasmid was incubated with purified Tn5 transposome (pre-loaded with Illumina adapters) at 55 °C for 10 min in the presence of 5 mM MgCl2 and 10 mM TAPS, resulting in an average fragment size of ∼400 bp. Reactions were treated with 0.5 volume of 0.1% sodium dodecyl sulfate for 5 min at room temperature and amplified with KAPA HiFi HotStart polymerase using primers containing 8 nt i7 and i5 index barcodes. Barcoded amplicons were purified with SPRIselect beads (New England Biolabs) and sequenced with a MiSeq kit (Illumina).

### Competent cell production

*E. coli* K-12 (ATCC 25404) was obtained from the American Type Culture Collection. Cells were cultured in ZymoBroth and made competent using a Mix & Go buffer kit (Zymo) according to the manufacturer’s recommended protocol.

### Phage plaque assays

*E. coli* harboring a candidate defense system or a pACYC184 empty vector were cultured in terrific broth at 37 °C in the presence of 25 μg/mL chloramphenicol. To 10 mL top agar (10 g/L tryptone, 5 g/L yeast extract, 10 g/L NaCl, 7 g/L agar) was added chloramphenicol (final concentration 25 μg/mL) and 0.5 mL *E. coli* culture, and the mixture was poured on 10 cm LB-agar plates containing 25 μg/mL chloramphenicol. Ten-fold dilutions of phages T2 (ATCC11303-B2) and T5 (ATCC11303-B5) in phosphate buffered saline were spotted onto the plates at 3 μL per spot. After overnight incubation at 37 °C, plates were photographed with a white backlight.

### Presence of UG/Abi RTs in Mobile Genetic Elements

Previously built HMM profiles for every UG/Abi group were used to retrieve sequences encoded in the PLSDB plasmid database (50). Firstly, ORFs were predicted for every nucleotide entry in the PLSDB database using Prodigal with default parameters and the -p meta option. After this, hmmsearch (E-value <1e-20) was used with every profile against the predicted ORFs. For every ORF, group assignment was done based on the top-scoring profile with at least >80% of sequence coverage. Sequences were aligned against the representative ones and manually inspected to remove false positives. The same procedure was employed against the mMGE and GPD databases (44,51). Further, phage prediction using the genomic neighborhood information was performed using the upstream and downstream nucleotide sequences (+-10kb) of every representative sequence as a query for the metasoftware WhatThePhage (52). Only sequences predicted to be encoded in phages/prophages by at least 3 different tools included in WhatThePhage were labeled as so.

### Analysis of viral genomes harboring UG27: annotation, taxonomy assignment, and host prediction

To search for UG27 systems in viral genomes, three different viral databases, IMG/VR (53), GPD (51), and mMGE (44) were used. First, searches with HMM profiles corresponding to UG27 RT, cluster 337, and cluster 346 were done using an E-value cut-off lower than 1e-20 against the viral proteins. Only genomes harboring RT, cluster 337, and cluster 346 were further considered, which rendered 3861 predicted viral genomes. Genomes were dereplicated at 95% sequence identity using dRep with default parameters as previously described (54), resulting in 1447 distinct genomes. In addition, dereplicated viral genomes were searched for Large Terminases (TerL) as a viral marker using the PFAM profiles PF03354, PF04466, PF03237, and PF05876 and a profile built from an alignment of TerL derived from a recent phylogenetic analysis (54). To identify these putative viral sequences and assign taxonomy, whole-genome comparisons and gene-content networks were made using novel phage families (54–56) as references. For whole-genome comparisons, fastANI (57) with parameter –fragLen 500 and default parameters were used. For gene-content networks, genes were called using a modified version of Prodigal (34) to allow for amber (TAG) or opal (TGA) stop codon reassignment (56) in those genomes with the corresponding suppressor tRNAs searched using tRNA-scan-SE (58), and/or an increase in the reassigned-coding density above 10%. After this, vContact2 (59) was used with the –db ‘None’ option and default parameter. To increase the resolution at the gene sharing level, an aggregate protein similarity tree was built as described earlier (55). To obtain family-level viral clades, the tree was rooted and cut at the levels reproducing the six proposed Crassvirales families (56). Finally, viral genomes harboring UG27 systems were annotated using Prokka (60) and the Phrog database (61) of viral orthologous proteins. GFF files produced by Prokka were then imported to R and depicted using gggenes (62) colored by Phrog functional categories (61). Host taxonomy prediction of UG27 viral genomes was done by performing CRISPR spacers comparisons against a reference database using CrisprOpenDB (63). As most of the taxonomy predictions pointed out towards bacteria present in the gut microbiome, viral genomes were compared against recent human gut metagenomic spacers database (64) and a custom collection of spacers found in CRISPR arrays from the human gut microbiome; compiled by running CRISPRCasFinder (65) on all metagenomic contigs from the HMP2-IBD database (66). For the comparison, BLASTn (67) with -task “blastn-short” option was used, and only hits with ≤2 mismatches and >95% sequence identity were kept.

### UG28 system RNAseq

E. coli K-12 (ATCC25404) containing a plasmid encoding RT (UG28) was grown to saturation at 37 °C in terrific broth in the presence of 25 μg/mL chloramphenicol. RNA was extracted using TRIzol (Thermo Fisher Scientific) and purified with a Direct-zol RNA MiniPrep Plus kit (Zymo). The purified RNA was then treated with 20 units of T4 polynucleotide kinase (NEB) for 3h at 37 °C. Following column purification, the sample was treated with 20 units of 5’ RNA polyphosphatase (Lucigen) for 30 min at 37°C. After an additional round of column purification, the RNA sample was used as input for an NEBNext Small RNA Library Prep for Illumina kit (NEB). Barcoded amplicons were sequenced on a MiSeq (Illumina) with 200 cycles for the forward read. Adaptors were trimmed using CutAdapt (68) with parameters --trim-n -q 20 -m 20 -a AGATCGGAAGAGCACACGTCTGAACTCCAGTCAC, and trimmed reads were aligned to the reference plasmid sequence using Geneious.

### Structured ncRNAs prediction

Paired RNAseq reads from SRX814863 were downloaded from the European Nucleotide Archive (www.ebi.ac.uk) and mapped to the RT (UG2) locus from *Klebsiella pneumoniae* ST23 (nucleotide accession number CP037742.1, corresponding to 100% nucleotide identity relative to the RNAseq reads) (Supplementary Figure S5) (69). Conserved structured ncRNA prediction was done on UG2 (DRT type 2) and UG3+UG8 (DRT type 3) UG/Abi groups. For UG2 members, nucleotide sequences 300 bp upstream of the RT were retrieved, whereas, for UG3+UG8, nucleotide sequences 300 bp downstream of UG8 RTs were retrieved. Using the information derived from the clustering and phylogenetic and taxonomic criteria, we divided UG2 and UG8 putative ncRNA sequences into subgroups. These subgroups were aligned using MAFFT-qinsi (28) and further examined in search of structure conservation in the neighborhood using CMfinder04 (70) and R-scape (71), as described previously (36). In groups of UG8 belonging to *Terrabacteria* phylum, no downstream structure or sequence conservation was observed, and predictions were made retrieving sequences 300 bp upstream instead, which showed broader structure and sequence conservation. Sequence logos for UG8 motifs were created using WebLogo 3 (72). For the prediction of structured RNAs in UG27 systems, the intergenic sequences located between UG27 RTs and cluster 336 were retrieved in representative sequences. Conserved RNAs structures were predicted on these sequences using CMfinder04 (70). Then, covariance models were built using cmbuild from the Infernal suite (73) and were compared against UG27 viral genomes described in the previous section using cmsearch from the same suite. The result of this search was then evaluated for statistically significant co-varying base pairs using R-scape (71).

## Results and discussion

### Expansion and classification of UG/Abi RTs

Recent phylogenetic analysis of prokaryotic reverse transcriptases revealed that UG/Abi RTs constituted a significant proportion (11%) of the whole RT landscape with 991 representative sequences (clustered at 85% of sequence identity) (5). By searching the NCBI public databases on complete or partial genomes and performing phylogenetic analyses of the most informative sequences (Methods), we could expand the number of UG/Abi RTs from 991 up to 5022 representative members, which represents a ∼5-fold increase compared to previously described UG/Abi RTs. Of these 5022 representative sequences, 325 belong to previously undescribed groups (Figure 1 and Supplementary Table S1).

**Figure 1.**
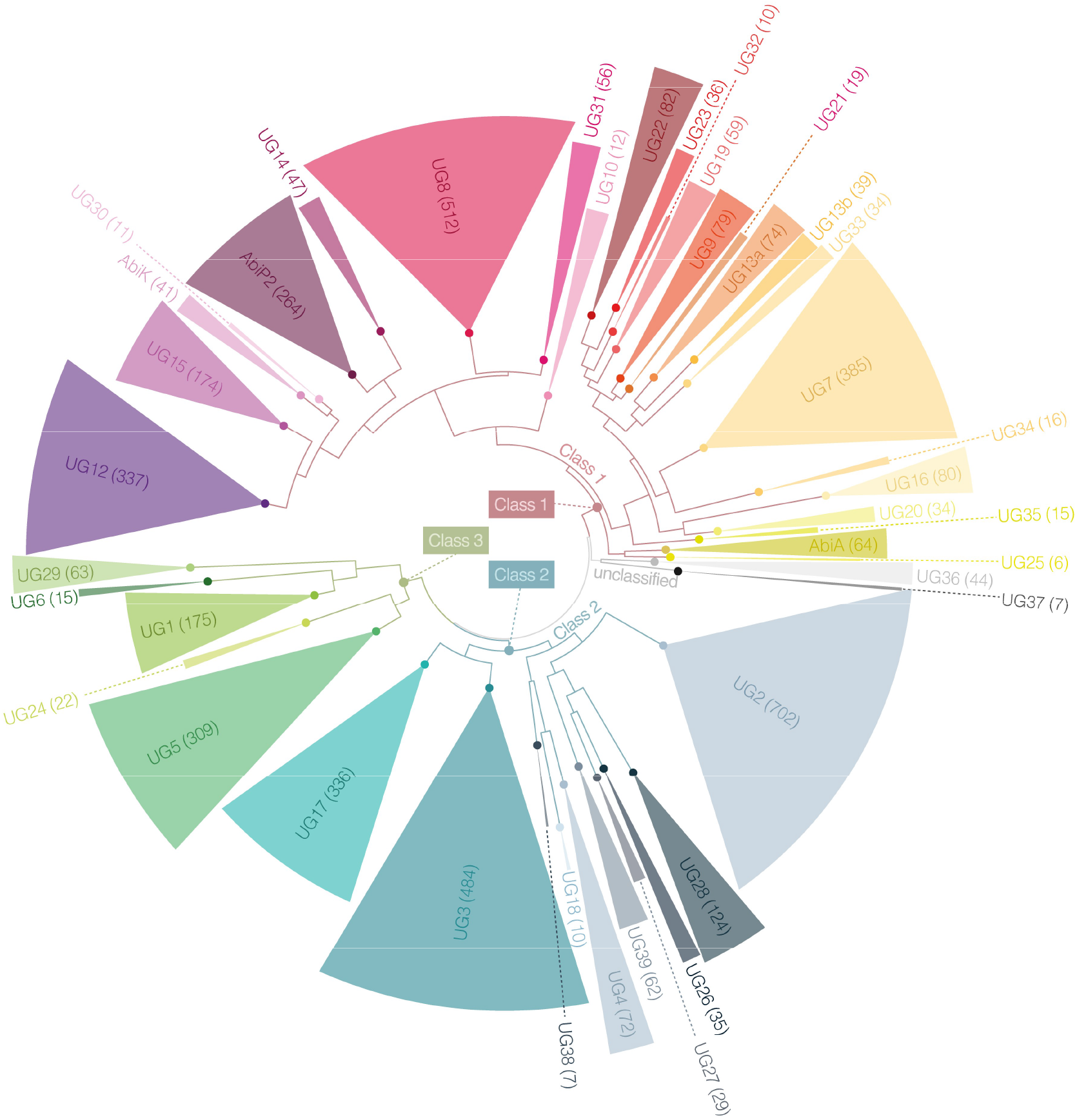
Phylogenetic tree of 5022 representative UG/Abi RT sequences. Alignment of the RT domains 1-7 was used as an input for FastTree (Methods). For visualization, midpoint rooting was applied using FigTree, and clades were collapsed to the UG/Abi group level according to the presence of reference sequences showing a FastTree support >0.70 (Supplementary File S1) with UFBoot and SH-aLRT support values in the IQ-TREE (not shown) >90%. Due to the presence of reference sequence in two groups, UG13 UG/Abi group was subdivided (UG13a and UG13b) according to phylogenetic criteria and protein length distribution. UG/Abi groups with no previously described reference sequences were considered to be novel and named starting from the last group discovered. Numbers in brackets and segments sizes represent the absolute number of representative sequences in each group. Branches corresponding o UG/Abi Classes 1, 2, and 3 are colored in brick red, turquoise, and olive, respectively. The phylogenetic tree can be found in Newick format as Supplementary File S1.

The representative dataset reveals that UG/Abi RTs predominate in Bacteria with only a few examples found in Archaea where they may have arrived by horizontal gene transfer. In Bacteria, these RTs are well distributed in Proteobacteria, Bacteroidetes, Actinobacteria, and Firmicutes as the main phyla with some groups mostly restricted to a particular phylum, (Supplementary Table S1 and Supplementary Figure S1) suggesting host-associated functional dependence.

To investigate the relationships between these novel sequences, we performed a phylogenetic reconstruction of the RT domain present in all UG/Abi RT representative sequences, increasing from 28 to 39 the number of well-supported UG RT groups and revealing the presence of three main clades (Figure 1). We then performed an exhaustive domain annotation of every UG/Abi group based on profile-profile searches and structural predictions, and a systematic analysis of the neighborhood of UG/Abi RTs in search of putative neighbor associated genes that were grouped into clusters (Figure 2 and Supplementary Table S2). This combined approach revealed that most of the UG/Abi RT groups (40 out of 42) are fused or functionally related to other genes or domains, thus disclosing a wide variety of genetic systems that were previously unnoticed. Besides, the annotation of domains, RT phylogeny, and prediction of associated genes made it possible to group UG/Abi RTs into three main classes described in the following sections (see Figures 1 and 2).

**Figure 2.**
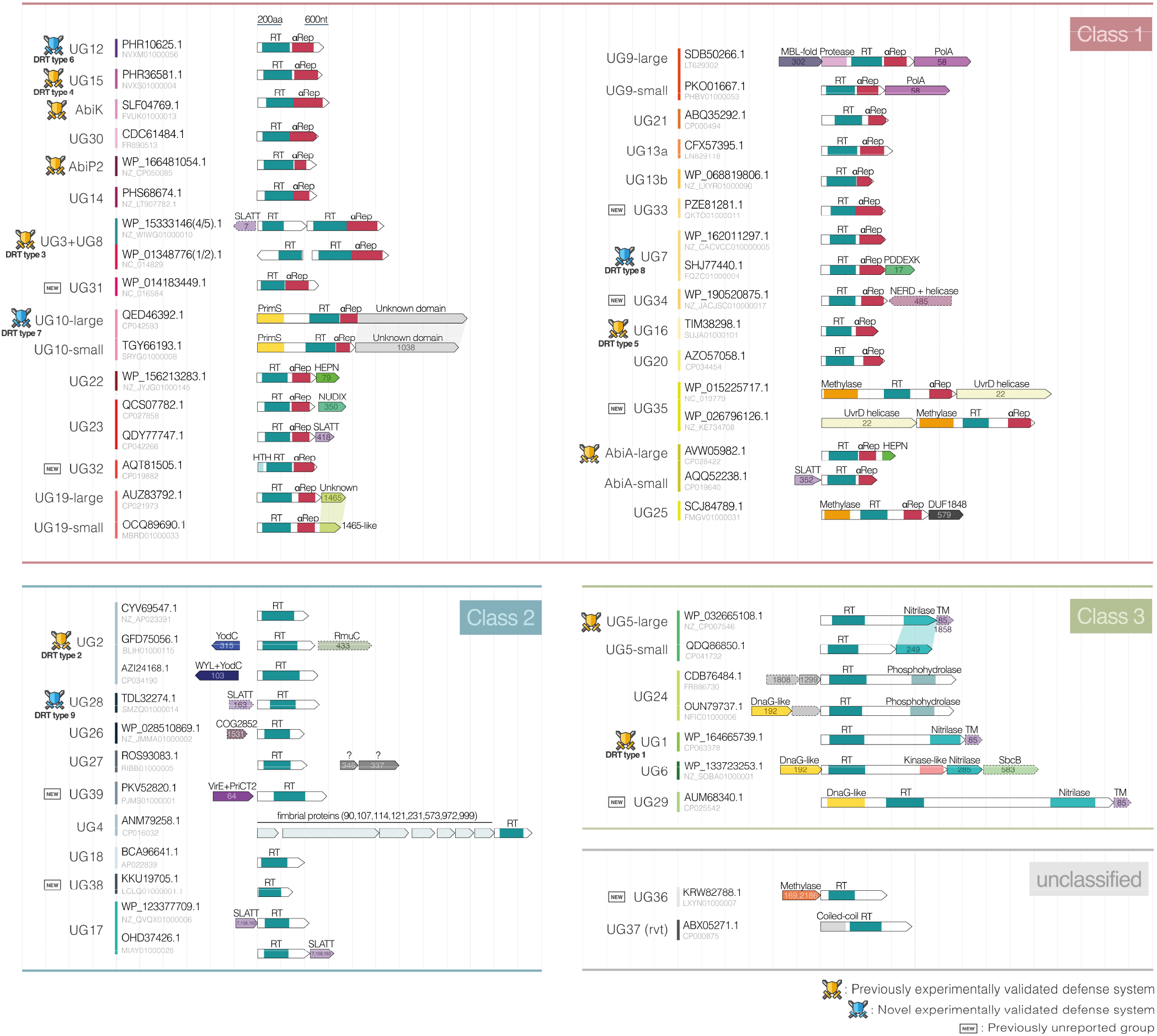
Graphical summary and classification of the various UG/Abi RT systems. Genes are represented as arrows of size proportional to their length. For clarity, the UG3+UG8 system is included within Class 1, but notice that the UG3 RT is phylogenetically placed within Class 2. The identified domains are represented in different colors. The NCBI Protein and Nucleotide accession IDs of chosen representative sequences are shown to the left. Numbers inside genes indicate the cluster to which they belong, and the names above indicate domain annotation. Genes with dotted lines represent genes that are occasionally present. Vertical dashed lines are represented to be 600 nucleotides (200 aminoacids) apart. Shadows between genes of different variants indicate similar domains. Icons on the left of each group’s name represent those that have been experimentally validated as defense systems and those that are considered to be novel.

### Class 1 UG/Abi RTs are fused to repeat-containing domains

Profile-profile searches of UG/Abi RTs against the COG database revealed that C-terminal regions of UG20, UG21, and UG23 members share remote homology with proteins containing HEAT-like repeats (Supplementary Table S2). To further investigate this feature, we performed structure predictions using trRosetta (45) and multiple sequence alignments (MSAs) of every UG/Abi RT group (see Methods). Interestingly, we detected alpha-helix repeats at the C-terminal regions of sequences belonging to a large clade comprising up to 24 UG/Abi RT groups, proposed to be denoted as Class 1 UG/Abi RTs. Despite the lack of overall detectable sequence similarity, structural alignments of those models highlighted a common structural motif consisting of a variable number of alpha-helix HEAT-like repeats (Figure 3) that will be named hereafter as αRep domain.

**Figure 3.**
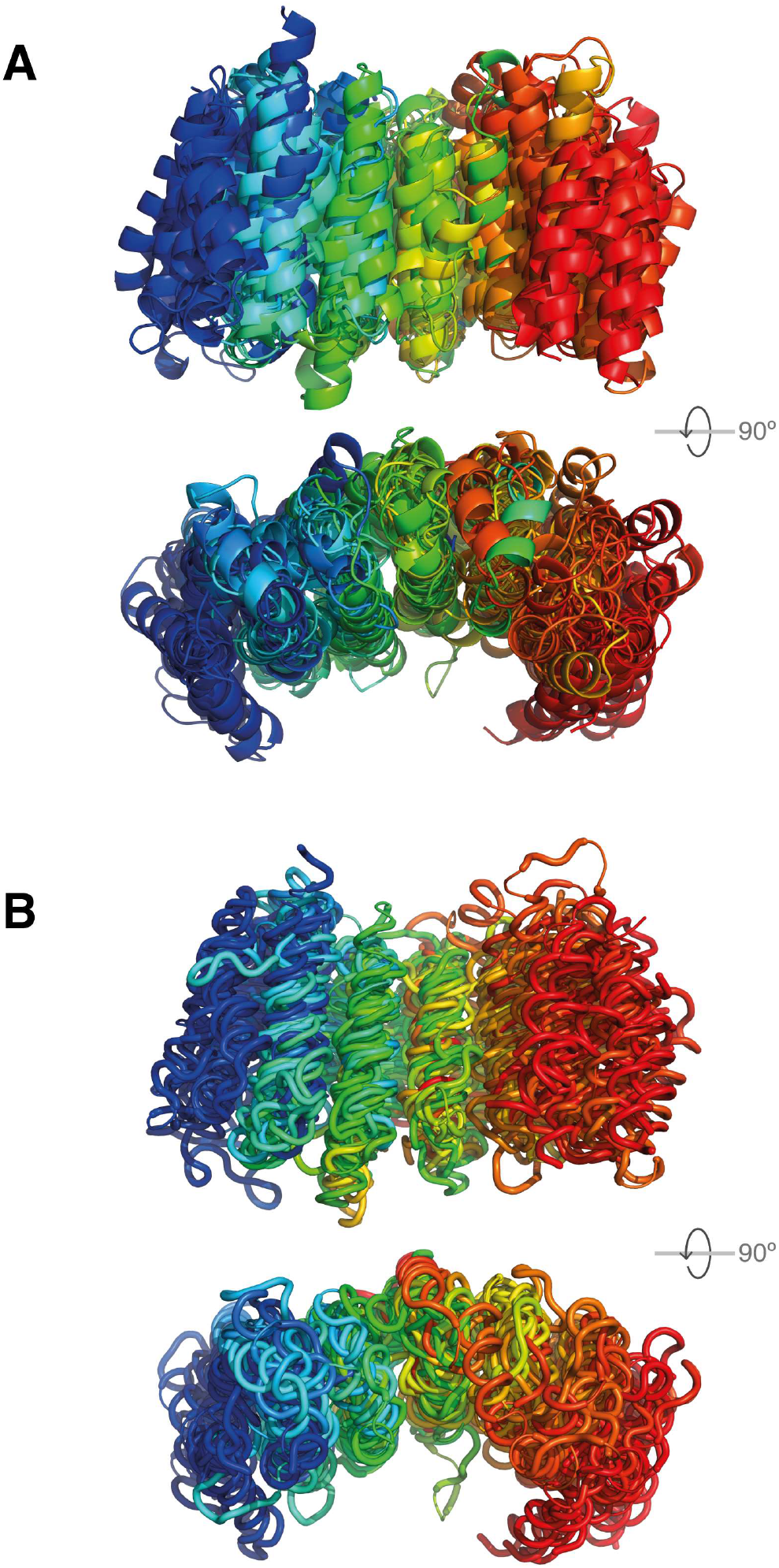
Multiple structural alignments of predicted protein structures of Class 1 UG/Abi RTs C-terminal regions visualized using PyMol2Rainbow rainbow spectrum A) Multiple structural alignments built using cealign feature from Pymol B) Multiple structural alignments built using mTM-align (Methods)

Further analyses revealed that other members of this Class are also fused to putative methylase (UG25, UG35), protease (UG9), primase (UG10), HTH (UG32), HEPN (AbiA), or unknown domains (UG10, UG19) (Supplementary Table S2), which points out a huge diversity in these systems, both at the mechanistic and biological level. Those fused domains can be sometimes encoded in a separate open reading frame (ORF) or absent, which allows a further division of some groups (UG10, UG19, UG9, and AbiA) into subgroups consisting of large (fused) or small variants (unfused) (Figure 2 and Methods). This phenomenon has been previously described for other RT-containing systems such as retrons, in which effector domains can be found both fused or adjacent to the RT (36). Genes in a separated ORF were also recovered by the gene-neighborhood systematic analysis (Methods), indicating that Class 1 UG/Abi RT groups constitute multi-domain systems, either encoded in a single ORF, or associated with other co-located genes.

The only characterized multi-gene system involving a Class 1 UG/Abi RTs is constituted by the association of UG8 with UG3, a Class 2 UG/Abi RT (see below, and Figures 1 and 2). These two genes constitute DRT type 3, which has been described to provide immunity against certain phages (21). Other uncharacterized multi-gene systems include genes with HEPN (UG22), NUDIX (UG23), SLATT (UG23, AbiA), UvrD helicase (UG35) domains, which are commonly found as part of prokaryotic defense systems such as CRISPR-Cas, retrons, RADAR (restriction by an adenosine deaminase acting on RNA), CBASS (Cyclic-oligonucleotide-based anti-phage signaling systems), Gabija, Abi or toxin-antitoxin pairs (18,21,22,36,74,75). On the other hand, as previously described, UG9 members are associated with a Family A DNA-polymerase (7), whereas a previously undescribed UG9 variant is associated with an MBL-fold metallohydrolase and a trypsin-like serine protease domain. Finally, the rest of the UG/Abi groups are also associated with unknown genes (UG10, UG25) or function as multi-domain stand-alone proteins, some of which (UG15, UG16, AbiK, AbiP2) have been described to provide immune functions (21,15).

#### Class 1 UG/Abi multi-gene systems including hydrolases

Some Class 1 UG/Abi groups are predicted to be associated or fused to enzymes that belong to the class of hydrolases (EC 3), such as nucleases, proteases, or helicases.

UG22 is predicted to be associated with neighbor protein cluster 79, which shares remote homology with HEPN domains (Figure 2 and Supplementary Table S2), a family of RNA-binding proteins that commonly function as endoribonucleases (76). This domain is also found at the C-terminal region of AbiA-large, which is thought to mediate immunity through phage RNA degradation or cell-dormancy induction. HEPN domains are commonly enriched in prokaryotic defense islands (77) and have been described as essential components of other defense systems/genes such as CRISPR-Cas, different Abi, and toxin-antitoxin systems, and ApeA (18,21). They also constitute type IX retrons together with a retron RT, a non-coding RNA, and a winged-helix domain-containing protein (36), which points out that RT and HEPN domains may have been co-opted several times across evolution to perform immune functions. Likewise, UG19 is associated with an unknown domain, either fused at the C-terminal region (UG19-large) or encoded in a co-located gene belonging to cluster 1465 (UG19-small) that shares very distant homology with HEPN domains (Supplementary Table 2). Other UG/Abi systems associated with nucleases include UG7 and UG34 (Figure 2). UG7 can be divided into two phylogeny-congruent variants. The first, less numerous variant is found to be associated with cluster 17, which shares remote homology with PD-(D/E)XK nucleases, a highly ubiquitous superfamily of nucleases related to Holliday junction resolvases (78). However, the second predominant does not appear associated with any gene, suggesting a likely accessory function of these nuclease-like proteins or the need for other nucleases that may act in trans. On the other hand, UG34 is more loosely associated with cluster 485, which contains a C-terminal helicase domain and an N-terminal nuclease-related (NERD) domain. In addition to their widespread distribution, both PD-(D/E)XK and NERD domains are enriched in prokaryotic defense islands (77). More specifically, PD-(D/E)XK-like domains are found in CRISPR-Cas and R-M systems nucleases, whereas it has been recently described that NERD nucleases can be found associated with prokaryotic Viperins (pVips) (79).

Other Class 1 UG/Abi groups are associated with hydrolases that are present in previously described defense systems. UG23 group members are predicted to be associated with cluster 350, which encodes a SMODS (Second Messenger Oligonucleotide or Dinucleotide Synthetase)-associated protein containing two transmembrane helices at the N-terminal region and a C-terminal NUDIX *(Nucleoside diphosphate linked* moiety X) hydrolase domain (Figure 2), which is thought to cleave nucleoside diphosphate molecules (80). This protein is known for being an effector protein of CBASS systems (75,80). Similar to Abi systems, CBASS possesses a sensor module that detects phage presence. In this case, it produces a cyclic-oligonucleotide signal and an effector module that subsequently recognizes this signal and triggers programmed cell death. Interestingly, in the case of AbiA and UG23, when the effector domain is absent (HEPN at the C-terminal region of AbiA-large or NUDIX hydrolase co-located with UG23) they are replaced by a SLATT protein, which is also sometimes co-located with the aforementioned UG3+UG8 system. The SLATT proteins contain 2-3 transmembrane helices and are predicted to function as pore-forming effector proteins associated with defense and conflict systems (80) and have been recently described to be part of novel defense systems such as RADAR or CBASS defense systems, in which their accessory role contributes to increasing the range of defense against different phages (21,22). These findings highlight the modularity of the different domains within these systems, being the sensor or effector domains easily replaceable by others with similar functions. Although associated with a wide variety of domains, the factors limiting the capacity of Class 1 UG/Abi RT and αRep domains to perform specific biological functions are yet to be explored, and it is possible that, although they are often associated with other genes or domains, these may have an accessory role, as in the case of the SLATT proteins in RADAR or DRT type 3 systems

#### Class 1 UG/Abi systems associated with transferases

Whereas some Class 1 UG/Abi multi-gene systems include hydrolases as putative effector proteins or domains, some other UG/Abi RTs are associated with enzymatic activities belonging to the class of transferases (EC 2), such as nucleotidyltransferases (DNA polymerases and DNA primases) or methyltransferases (also known as methylases).

UG9 RTs function together with a Family-A DNA polymerase (PolA, cluster 58) (7) and can be divided into two different variants. The first, less frequent variant, is associated with hydrolases, with the RT being fused to an N-terminal putative trypsin-like serine protease and an MBL-fold metallohydrolase found upstream (Figure 2). This architecture is reminiscent of Avs1 defense systems, in which the protease is fused at the N-terminal region of a protein containing a STAND ATPase domain and the MBL gene is also located upstream (21). The second UG9 variant is not associated with an MBL-fold metallohydrolase and the RT does not harbor an N-terminal protease domain, thus pointing out that both the metallohydrolase and protease domains would operate together. In this way, the MBL/protease pair may function as an accessory or effector module of different defense systems, thus being frequently exchanged among different antiphage systems in a similar way to NUDIX hydrolases or SLATT proteins.

UG10 RTs are very large proteins (>800aa) that also associate with replication-related domains, as they harbor an N-terminal PriS domain (COG4951) that belongs to the AEP (archaeo-eukaryotic primase) superfamily (81,82). The fusion of AEP primases to RTs has been previously described as part of defense systems such as CRISPR-Cas (5) or Class 3 UG/Abi member DRT type 1 (21) and suggests that similar to HEPN domains, both the RT and primase domains may have been co-opted together several times across evolution to perform immune functions. In addition to the N-terminal PriS domain, UG10 members are associated with an unknown domain with ∼800aa that can be either fused to the C-terminal region (UG10-large) or encoded in a separate CDS that belongs to cluster 1038 (UG10-small).

Finally, both UG35 and UG25 RTs harbor an N-terminal methyltransferase domain. To our knowledge, this is the first reported case of a predicted methylase-RT fusion in prokaryotes, although the association of methylase and other bacterial RTs has been previously described (87). Apart from being fused to the methylase domain, UG35 RTs are associated with cluster 22, which is identified as a putative UvrD-like helicase. However, UG25 members are not associated with a helicase but with cluster 579, a cluster containing a DUF1848 domain of unknown function.

### Class 2 UG/Abi RTs do not harbor any additional known domain

While Class 1 UG/Abi RTs are fused to αRep and other known domains, Class 2 UG/Abi RTs do not bear any additional known domain. Instead, the 9 different groups belonging to this class harbor the RT domain and are sometimes associated with other gene(s) with DNA-binding or DNA-modifying activities that seem to play an accessory role. Although their C-terminal domains do not appear to have a conserved sequence or predicted structure, further genetics, and structural investigations can make it possible to reveal their possible shared characteristics.

UG2 is the largest group and it can be classified into three different variants (Figure 2). The first variant is a single-gene UG2 RT, whereas the second and third variants are co-located with genes (clusters 315 and 103, respectively, see Supplementary Table S2) that share an N-terminal YodC (COG5475) domain. Besides YodC, cluster 103 presents a WYL-like domain at the C-terminus. Although very little is known about YodC domains, the WYL domain is known to be a group of diverse transcription factors that can regulate a response when binding to RNA molecules (83). This domain is enriched in defense islands (21) and it has been proved to regulate some CRISPR/Cas systems and other defense systems such as the abortive infection AbiG system (83,84). Some of the second variant members have another associated cluster in the neighborhood (cluster 433 in Supplementary Table S2), which contains an N-terminal TM (transmembrane) domain followed by a COG1322 domain (RmuC DNA anti-recombination protein) with a restriction endonuclease-like fold and coiled-coil regions (78) (Supplementary Table S2). For UG2, it has been demonstrated that the sole presence of the RT domain (along with an ncRNA) is sufficient to confer anti-phage functions, thus suggesting that in this case, the associated clusters may have an accessory role.

Other UG/Abi systems possibly associated with accessory proteins include UG28 and UG26. UG28 is sometimes associated with cluster 163 which is identified as a SLATT protein, whereas UG26 can be co-located with cluster 1531, which is a COG2852 domain-containing protein annotated as YcjD, a Very-short-patch-repair endonuclease (Figure 2 and Supplementary Table S2) commonly associated with DNA repair mechanisms and linked to PD-(D/E)XK nucleases (19). RTs belonging to the UG39 group are predicted to be associated with the protein cluster 64 (Figure 2) which contains an N-terminal VirE domain and a C-terminal PriCT-2 domain (Supplementary Table S2). Both domains are commonly found in PrimPol proteins (primase-polymerases) from the AEP superfamily (81,82).

UG4 RTs were previously described to be fused to fimbrial domains (6). In this case, we found a subgroup of proteins from the UG4 group widely associated with cluster 90, cluster 107, cluster 114, cluster 121, cluster 231, and cluster 573 which are predicted to be fimbrial proteins (Figure 2). However, we were unable to find fimbrial domains fused to any of the representative RTs classified as UG4 in our dataset, suggesting that previous identification of the fimbrial domain in UG4 RTs may be due to an inaccurate prediction of the ORF boundaries (6).

RTs from the UG27 group are putatively associated with clusters 346 and 337. Profile-profile searches revealed that none of these clusters showed homology to known domains. This suggests that they may represent novel gene families with unknown functions. The same happens with UG18, which is sometimes co-located with cluster 2007 that does not harbor any identifiable domain. Nevertheless, the validity of UG18 RTs, similar to UG38, cannot be fully tested, since both of them are small and infrequent groups present in partial genomes from metagenomic sources. Whereas UG18 is harbored by *Gammaproteobacteria sp*ecies (Supplementary Table S1) UG38 members are present in *Parcubacteria*, *Nealsonbacteria*, and *Atribacteria* species, which belong to the Candidate Phyla Radiation (CPR), a recently described group of mostly unculturable nanobacteria that constitute a novel lineage with extremely reduced genomic repertoires due to their predicted symbiotic and/or parasitic lifestyles (85). Because of the nature of their genomic sources, both UG18 and UG38 UG/Abi groups could be significantly expanded by performing exhaustive metagenomic searches.

Finally, reverse transcriptases belonging to UG17 are strongly associated with clusters 7, 163, and 158 which are identified as SLATT (SMODS and SLOG-associating 2TM) proteins. Experimental approaches have described that, under certain conditions, UG17 systems influence SbcB (exodeoxyribonuclease I) essentiality in some *E. coli* strains, as an insertion within the RT abrogates SbcB essentiality (86). This implies that the presence of UG17 makes SbcB essential, thus denoting that UG17 systems may have a toxic effect possibly mediated by the SLATT gene. Similar to Class 1 UG/Abi RTs, different Class 2 UG/Abi RT groups are associated with SLATT proteins. As we have described, these proteins are present in different scenarios, either having an accessory role (UG3+UG8, UG28, RADAR, CBASS), replacing effector proteins (AbiA, UG23), or strongly associating with the RT (UG17). In a context of high evolutionary pressure and gene turnover, the gap left behind by outgoing effector modules may be filled with the recruitment of highly mobile and adaptable SLATT proteins that can eventually lead to a long-lasting stable association such as UG17.

### Class 3 UG/Abi RTs are associated with (phospho)hydrolase domains

Finally, Class 3 UG/Abi RTs are large proteins (generally larger than 1000 aa) with an RT domain associated or fused to C-N hydrolase (also known as nitrilase) or phosphohydrolase domains. UG1, UG5, and UG29 RTs share a C-terminal C-N hydrolase domain. In the case of UG1 (1200-1300 aa), the characteristic C-N hydrolase domain is thought to be essential for their function, as mutations in this domain abrogate immunity against phages provided by DRT type 1 in *Klebsiella pneumoniae* (21). In addition, UG1 can be frequently found to be associated with cluster 85, which is identified as a transmembrane (TM) protein, and UG5 can be found in two different variants, the first of which (UG5-large) shares a similar domain architecture with UG1 with a fused C-terminal C-N hydrolase domain and a TM protein (cluster 1858, Supplementary Table S2) sometimes encoded downstream. However, the second variant (UG5-small) presents the C-N hydrolase domain encoded in a separate ORF (cluster 249) and it is not associated with a TM protein.

UG29 RTs are by far the largest proteins in our dataset, with almost 2500 amino acids. These proteins display a characteristic DnaG-type primase at the N-terminus, followed by an RT domain, and a C-terminal hydrolase domain. Although encoded in different CDS, UG6 presents the same hydrolase and DnaG-type primase domains in associated clusters 285 and 192 (respectively), thus indicating a possible common mechanism of action. However, in this case, UG6 also displays an unknown C-terminal domain that shares remote homology (Supplementary Table S2) with kinase domains but lacks the characteristic catalytic amino acids. DnaG-type primases, which contains a CHC2-type zinc-finger followed by a TOPRIM (Topoisomerase-primase) domain, were traditionally thought to be only involved in bacterial chromosome replication whereas AEP primases such as that found in Class 1 UG10 were thought to mediate the same mechanism in archaeal and eukaryotic species (82). Although functionally related, both the AEP and DnaG-type primases are evolutionarily and structurally distinct, revealing that the association of UG10 RTs to AEP primase is an event independent from the acquisition of DnaG-type primases by Class 3 UG/Abi RTs. This similarity of domains explains why members of UG29 have been previously mislabeled as UG10 (21).

On the other hand, UG24 members encode a predicted C-terminal phosphohydrolase domain (COG3294) instead of the above-mentioned hydrolase domain, thus pointing out that the enzymatic reaction that gives rise to the immunity could be carried out indistinctly by C-N hydrolase or phosphohydrolase domains. Furthermore, depending on the genes to which it is associated, UG24 can be divided into two variants, one of them associated with cluster 192 (the same DnaG-type primase found to be associated with UG6) and one of them associated with two genes of unknown function (clusters 1808, 1299).

Although the C-N hydrolase family generally performs cellular functions related to nucleotide metabolism and small molecule biosynthesis, the hydrolases found in Class 3 appear to be more closely related to each other and form a clade distinct from the other nitrilases, leaving open the possibility that their biological function is completely different and may be more related to immune functions (21). In addition to the hydrolase, phosphohydrolase, and primase domains, all these proteins share an extension of amino acids generally located between the RT domain and the respective C-terminal domains with unknown function. This domain could provide these large RTs with sufficient molecular flexibility to orchestrate the different enzymatic reactions catalyzed by their different domains and the associated genes.

### Unclassified UG/Abi RTs

Other RTs belonging to the UG/Abi RT family, namely UG36 and UG37, remain unclassified due to the absence of characteristic domains or their phylogenetic positioning, although future expansions in number and diversity may clarify their classification. In the case of UG36, we found that it is associated with genes that present a methylase domain grouped indistinctly in clusters 169 or 2186 (Supplementary Table S2). The domain architecture of this system is reminiscent of UG25 and UG35 RTs, in which the methylase domain is fused to the N-terminal end of the RT-containing protein. Thus, UG36 and UG25/UG35 systems may share a common ancestor. However, there is no evidence that they present a C-terminal αRep or similar domain. Similar to the associations of HEPN or primase with RTs, the methylase domain has been associated with an RT at different points in evolution. A less tight link between RTs and methylases appears to occur in *Lactococcus lactis*, in which rRNA methylation acts as a mechanism to prevent retrotransposition of endogenous group II introns (87).

On the other side, UG37 corresponds to *rvt* elements. The *rvt* are single-copy genes of unknown function that, in addition to the RT domain, contain an N-terminal coiled-coil domain that is responsible for its multimerization *in vitro* (88). They can be found in all eukaryotic kingdoms and a few bacterial genomes and, although they show a patchy distribution, they are not components of retrotransposons or viruses (89). *In vitro* studies suggest that *rvt* are the first chromosomal RTs described to have a protein-priming DNA synthesis initiation mechanism, which is thought to be dependent on the unknown C-terminal domain and the RT catalytic domain itself (88). Protein-priming has been described also for Class 1 AbiK, where the C-terminal domain has been also hypothesized to play this role and it is essential for abortive phage infection (14).

### High occurrence of UG/Abi RTs in Defense Islands

As we highlighted in the above sections, it has been previously shown that different RT families are involved in defense against phages (4). Recently, some UG/Abi RT groups have been demonstrated to function as defense systems named as DRTs and are located within defense islands (21,77). Besides AbiK, AbiA, AbiP2, and DRTs, it was previously shown that UG1, UG5-large, UG29 (named as UG10 in (21)), UG7, and UG9 are also enriched in defense islands but their immune function has not been experimentally validated (21). Interestingly, a recent preprint (90) has revealed that P2-like and P4-like prophages possess variable genomic regions encoding previously known anti-phage systems and 14 novel characterized defense systems, including a UG5-large system found in *E. coli E101*. Within these variable genomic regions, we also found, in addition to UG5-large, type II-A1 and II-A2 retrons (36) and RTs from UG/Abi groups such as UG7, UG3+UG8 (DRT type 3), UG15 (DRT type 4) and UG17 (Supplementary Table S4), thus suggesting that UG7 and UG17 may also provide immune functions.

In the case of UG5-large, the associated cluster 1858 (TM protein) is necessary for the system to perform immune functions (90). However, previous attempts to describe these immune functions did not take into account this protein and the same may occur in other groups in which associated genes were overlooked. Particularly, this seems to occur also in the case of UG7, in which the tested sequence falls into the variant that is associated with a PD-(D/E)XK nuclease (cluster 17).

Given that many of the UG/Abi RT groups have members that have been validated as defense systems or have been described to be enriched in defense islands, we were interested to know if all UG/Abi RTs are predicted defense genes. To investigate their defense association, we searched for the presence of previously reported defense genes in the vicinity of homologs from each RT group (Methods). We observed that the majority of UG/Abi RT groups are indeed enriched in known or predicted defense islands and are likely to perform immune functions (Supplementary Table 3). Of the 42 UG/Abi RT groups, members of at least ten groups (AbiA, AbiK, AbiP2, UG1, UG2, UG3, UG5, UG8, UG15, and UG16) have been experimentally shown to confer anti-phage defense activity (13-17,21,90), consistent with all of them having high defense association frequencies. Seven other groups (UG7, UG9, UG12, UG14, UG17, UG28, and UG29) also had strong defense enrichment, with 0.15 association frequency or greater across a minimum of 50 homologs; therefore, they are predicted as novel defense genes (Supplementary Table 3). The remaining groups had either somewhat lower defense association frequencies or fewer than 50 homologs. For these groups, individual genomic loci were examined for instances with particularly strong defense association signatures (Methods and Supplementary Figure S2). Based on this evidence 13 additional groups (UG6, UG10, UG13a, UG13b, UG19, UG21, UG22, UG24, UG25, UG30, UG31, UG35, and UG36) were predicted to have a defense function.

No defense enrichment was detected for UG4 and UG27, consistent with their predicted non-defense function (see below section). On the other hand, defense predictions were inconclusive for the remaining 10 groups (UG18, UG20, UG23, UG26, UG32, UG33, UG34, UG37, UG38, and UG39), thus requiring further investigation (e.g., search for more homologs or experimental observation).

To confirm that RTs with a high defense score were involved in defense, we heterologously reconstituted some candidate systems (UG10, UG12, UG28, and UG7, and UG15 as a positive control) in *E. coli* K-12 and challenged them with T2 and T5 coliphages following a similar procedure to that previously described (21). Phage sensitivity of the RT-containing bacteria was compared to that of an empty vector control vector by performing phage plaque assays (Figure 4 and Methods). We observed anti-phage activity for UG10, UG12, UG7, and UG28. In particular, UG10 and UG28 were found to be quite active against T2 phage, whereas UG7 and UG12 were active against T5 phage. Following the previous nomenclature for UG/Abi RTs involved in defense (21), we propose to name UG12, UG10, UG7, and UG28 as DRT type 6, 7, 8, and 9 respectively (Figure 2). For Class 1 UG/Abi RTs, in addition, we found that the C-terminal αRep domain is required for immunity, as deletion of a small portion (38aa) from the C-terminus of UG15 abrogated immunity (Figure 4).

**Figure 4.**
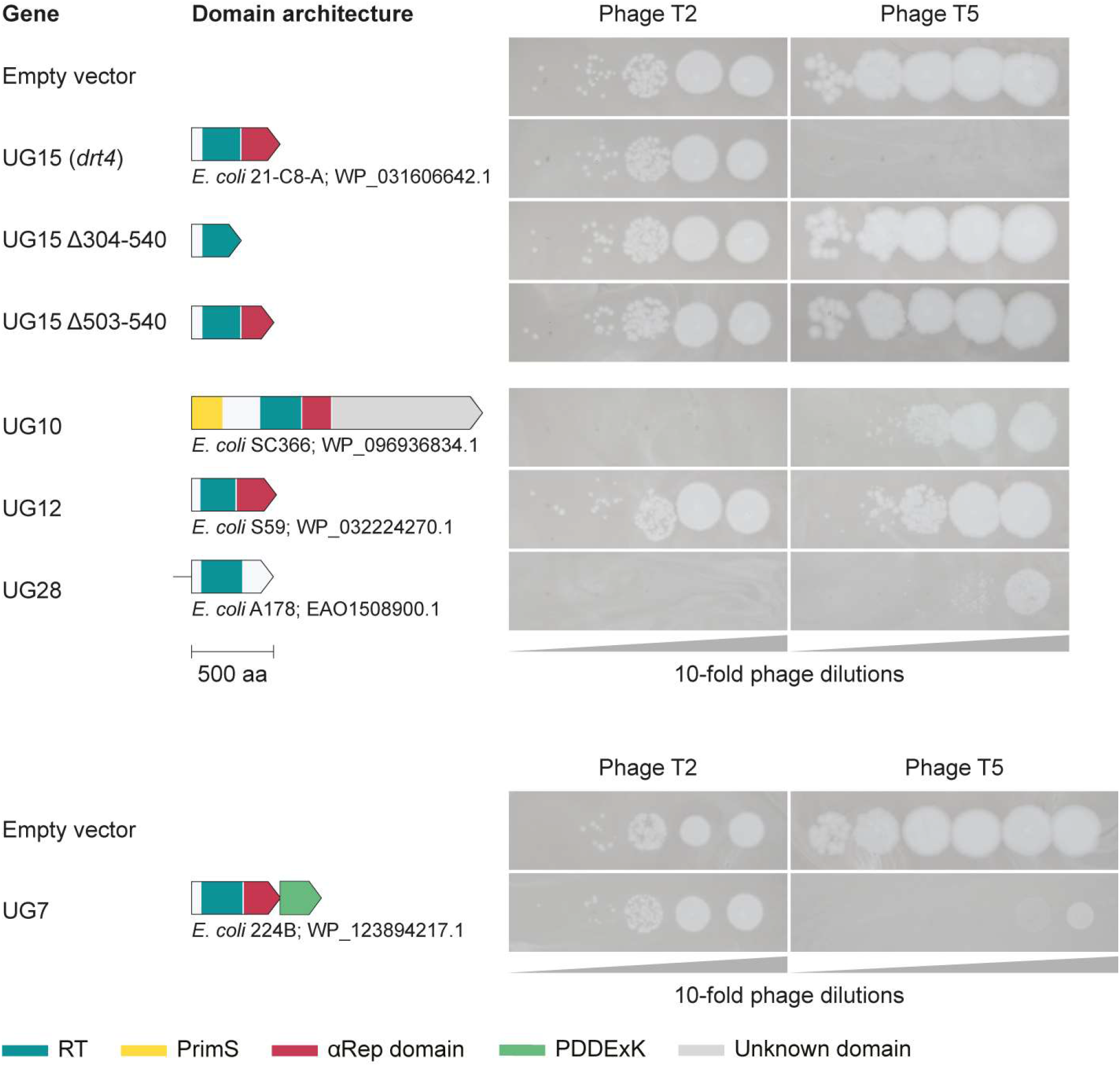
Phage plaque assays showing resistance of *E. coli* harboring plasmids with different UG/Abi RTs against coliphages T2 and T5. UG15 variants containing partial and complete deletions of the C-terminal αRep domain are also included. RT, reverse transcriptase; PrimS, primase.

These results reveal that the UG/Abi RTs lineage constitutes a family of RTs largely involved in defense against viruses, as previously highlighted for other prokaryotic RT families such as retrons (21–24). In addition, the enormous diversity of domain architecture and associated genes suggests that the RT domain activity might require other enzymatic activities or domains to play an immune role, as we have demonstrated for Class 1 αRep domains.

### Presence of UG/Abi RTs in Mobile Genetic Elements and Viral genomes

It has been previously described that some UG/Abi RTs can be present in MGEs, including plasmids (AbiK and AbiA), prophages (AbiP2 and other groups in P2-like prophages), among others (17,90,91). In addition, UG15 members have been also found within the *Helicobacter pylori* accessory genome (HPSJM_07740 gene of *H. pylori* SJM180) (92). To find which UG/Abi RTs were encoded in MGEs, we analyzed the genomic context of every representative sequence in search of phage signatures using the metasoftware WhatThePhage (52) and searched MGE databases (50) with all prokaryotic RTs HMM profiles (Methods).

As a result, we obtained that some UG/Abi groups tend to localize in putative MGEs. Particularly notable is the case of UG27, most of whose members are classified as viral by WhatThePhage and whose presence in the GPD and mMGE databases is remarkable (44,51) (Supplementary Table S4). Other groups such as AbiP2, UG2, UG3+UG8, UG5, UG7, UG12, UG15, and UG17 had several representative sequences predicted to have viral origin according to WhatThePhage. Of them, AbiP2 and UG2 were found to be frequently encoded in plasmids too, whereas AbiK and UG4 were mainly present only in plasmids. This suggests that AbiP2 and UG2 are highly mobile across different MGEs, and points out that the lack of detection of UG4 and UG27 in defense islands may be due to the fact that they are located in plasmids and viruses, respectively. Both may have been recruited by MGEs due to their shared evolutionary dynamics (93) and may perform a distinct biological role, such as acting as an addiction module or as an anti-defense system.

To verify the presence of UG27 in viral genomes, we expanded the number of UG27 systems by searching on IMG/VR, GPD, and mMGE databases (44,51,53) obtaining 1447 dereplicated genomes harboring the complete system (781 of which also encode a TerL homolog, Supplementary Figure S3). In an attempt to assign taxonomy to these viral sequences or test whether they belong to a known family, we compared them against reference databases and a compendium of new phage families (54–56) using ANI (average nucleotide identity) and gene-content network information (Methods). At the ANI level, some UG27-containing viral genomes share similarities with recently proposed orders of bacterial viruses, including *Crassvirales* and *Friedlandervirales* (55) and viruses from the proposed *Quimbyviridae* family (54) (Supplementary Table S5). Most of the UG27-containing genomes however showed very distant to zero similarity to any currently classified bacteriophage family even at the protein level, so we constructed an aggregate protein similarity (APS) tree (55) to determine whether the phages belonged to known or novel viral families. After cutting the APS tree at the viral family level, by using the six proposed *Crassvirales* families (56) as references, some of the UG27 containing phages co-clustered with *Quimbyviridae* (54) and various families within the proposed *Crassvirales*, *Twortvirales* and *Freidlandervirales* viral orders (55) (Supplementary Table S6). Four main clusters (205, 206, 255, and 265) accounted for 1350 out of 1447 UG27-containing genomes. Of these, family cluster 255 was co-clustered with genomes belonging to diverse families within the Friedlandervirales order, whereas family cluster 265 was co-clustered with members of the *Sylversterivirdae* family belonging to the *Crassvirales* order. On the other hand, half of the UG27-containing phage genomes were found within three separate family-level clusters (205, 206, and 291) that likely represent novel viral families (Figure 5). The largest of the clusters, with 444 members was more expansive than any other known dsDNA viral family data, whereas the second largest, at 252 members was around the same size as the largest crass-like family *δ-crassviridae*. The gene content of those clusters (Supplementary Figures S4 and S5) was characteristic of *Caudoviricetes* class head-tail bacteriophages. Despite the predominance of these two family-level clades in gut metagenomic data worldwide, the families appear to have gone unnoticed so far, and we propose naming them *Astarteviridae* and *Habisviridae*, respectively. (Families 206 and 205 in Figure 5, Supplementary Table S6 and Supplementary Figures S4 and S5).

**Figure 5.**
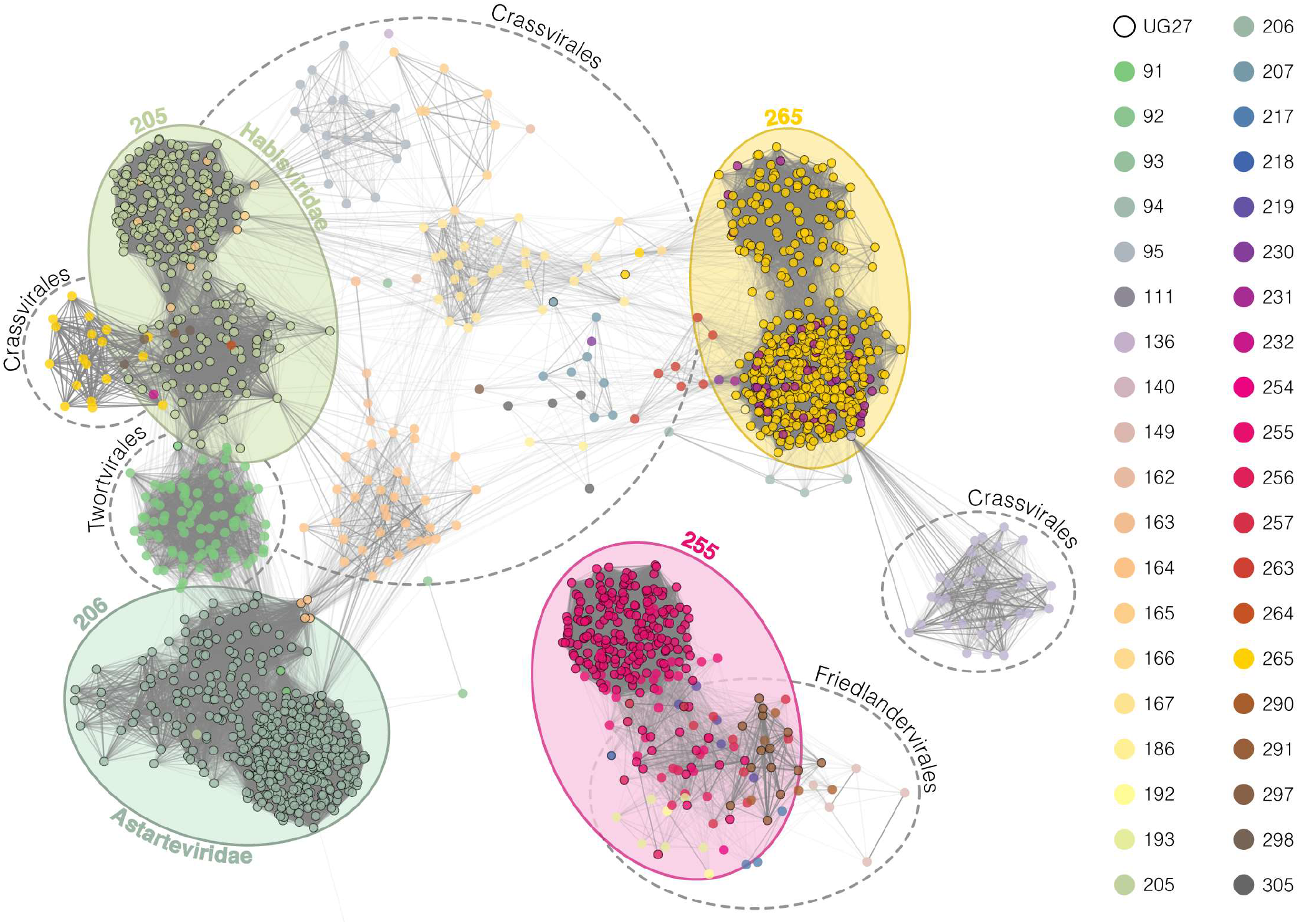
Gene-sharing network of UG27-containing viral genomes dereplicated at 95% and similar viruses found in reference databases (Methods). Nodes represent viral genomes and edges connect genomes with similar gene content. Edge opacity is proportional to vContact2 weights which represent the significance of the relationships. Node colors indicate the family to which every genome belongs based on the APS-tree, and circle color shades around nodes represent the four major families found. Nodes with no connections to UG27-containing nodes were removed to improve the visualization.

According to spacer-based host taxonomy assignment and viral genomes metadata (Methods), we obtained that most of the predicted hosts of these viral genomes belong to the Bacteroidetes and Firmicutes phyla (Supplementary Table S7), within species highly prevalent in human gut microbiomes, which could be influenced by the origin of the data. Next, we tried to infer whether these putative phages were predicted to be active or not by comparing them with databases of CRISPR spacers derived from human gut metagenomes. To this end, we used a publicly available collection of spacers extracted from 11, 817 human gut metagenome datasets (64) and an in-house spacer database that was built by running CRISPRCasFinder (65) on the Integrative Human Microbiome Project – Inflammatory Bowel Disease metagenomic dataset (66) (Methods). By doing this, we obtained that 1346 out of 1447 dereplicated viral genomes harboring UG27 (93, 02%) were targeted by at least 1 spacer, with 1197 (82%) being targeted by 5 or more spacers, suggesting a recent active role of these viral genomes in their natural environment. We also noticed that, in some cases, spacers targeted the intergenic sequence located between UG27 RT and the associated cluster 336, highlighting a possible functional role of this non-coding region (Supplementary Figure S6).

### Group-specific structure of ncRNAs in DRTs and UG27

The ncRNA present at the 3’ end of the UG3+UG8 system has been described to be essential for immune functions (21). Since most UG2 and UG28 homologs are associated with a sizable predicted non-coding sequence in the genomic DNA, located at the 5’ end of the RT ORF, we hypothesized that these sequences also encode for an ncRNA. To investigate the presence of UG2 and UG28-associated ncRNAs, we paired RNAseq reads to UG2 and UG28 (Methods) systems. This data revealed highly-expressed ncRNA at the 5’ end of both locus with read coverage >1000-fold greater than that of their respective RT mRNA (Supplementary Figure S7)

After this, we predicted conserved secondary structures in the vicinity of UG2, UG8 and UG28 using a methodology similar to the one previously used to uncover some ncRNA present in retrons (36). Due to low sequence and structure conservation, UG2 and UG8 upstream/downstream sequences were grouped into 13 and 20 groups, respectively, according to the RT phylogeny and taxonomic criteria (Supplementary Figures S8 and S9). For UG28, however, the high degree of conservation allowed all sequences to be grouped into a single cluster. Then, consensus RNA structure predictions were made on every group.

Overall, some UG2 upstream sequence groups showed evident secondary structure conservation, whereas we were unable to detect consensus structures in some others, possibly due to a great divergence both at the sequence and the structure level and the low number of homologs (Supplementary Figures S8 and S10). Generally, the predicted secondary structures in predicted UG2 ncRNA greatly vary among the different groups, and there are no clear signs of inter-group sequence conservation, thus suggesting that the interaction between the UG2 RTs and the predicted ncRNAs may be highly specific.

On the other hand, the RNA structure conservation is clearer in UG8, yet maintains a high divergence at the sequence level. Furthermore, there is a clear distinction into two groups within UG8; those with the canonical UG3-UG8-ncRNA architecture in Proteobacteria and Firmicutes, and those present in Terrabacteria, which have a different architecture (UG3 and UG8 frequently encoded in opposite strands and the ncRNA upstream of UG8) (Supplementary Figure S9). In the first case (groups 1-17), the existence of a “motif” that could be related to the function of the RNA is evident. This “CACACA”-like motif is globally conserved among these groups and it seems to be exposed in the RNA structure predictions, but the distance to the end of the RT varies greatly (Supplementary Figures S11 and S12). We speculate that this motif could be a recognition/attachment motif or a binding site that promotes the association of the UG8 RT. Although all of them share this motif, there are no signs of inter-group sequence similarity in other regions, highlighting that the importance of this possible ncRNA may be more dependent on its structure than in its sequence.

In contrast to the ncRNAs in UG2 and UG8, conservation at both the structural and sequence level in the predicted ncRNAs for UG28 was surprisingly high (Supplementary Figure S13), suggesting that the response mechanism of this particular system might be much more conserved and nonspecific.

Furthermore, we also analyzed the intergenic region located between UG27 RTs and cluster 336 in search of structurally-conserved ncRNAs revealing the presence of a putative conserved ncRNA (Supplementary Figure S6) with three long stem-loops that may be relevant for the functioning of UG27 systems within viral genomes. Due to the nature of the data used for the prediction (mostly coming from metagenomic sources and viral genomes predictions), the existence of this ncRNA should be validated experimentally, although it highlights its importance and the possible requirement for such a system to be functional.

The presence of ncRNAs in Class 1 (UG3+UG8) and Class 2 (UG2 and UG27) UG/Abi systems highlights the possible existence of non-coding RNAs in other groups of UG/Abi groups. In the case of UG3+UG8 (DRT type 3), ncRNA plays an indispensable role in achieving immunity against viruses, suggesting that their specific interaction with the RT may be tightly regulated and is responsible for triggering a signal or generating modifications such that the host can cope with a bacteriophage invasion by a yet unknown biological mechanism.

## Conclusion

Through computational methodologies, in this work we have proposed the existence of a new family of RTs, hereafter proposed to be named UG/Abi RTs. By expanding the number and diversity of these RTs, we attempted to clarify their relationships and confirm their evolutionary and biological similarities. In addition, we built a phylogeny-congruent categorization of UG/Abi RTs in three major Classes, and we have experimentally demonstrated the defensive function of four new groups grouped into Class 1 and Class 2 UG/Abi RTs.

In the context of UG/Abi RTs, the phylogeny of the RTs correlates well with the presence/absence of fused or associated modules, which suggest some degree of co-evolution and a functional limitation to operate with non-specific modules. That notwithstanding, in the highly dynamic evolutionary context of defense systems, these barriers are easily overcome and exchanges of modules/genes between different systems can be observed. The promiscuity of UG/Abi RTs in associating with different effector modules reveal their vast diversity as well as the plasticity of these systems, which likely expands their target molecules and anti-phage actions, possibly compatible with other defense systems, acting along and/or in coordination with some of them, as previously reported for some retrons that serve as a “second line of defense” (22). However, even though different RT families (retrons, CRISPR-associated RTs, UG/Abi RTs) associate with the same type of proteins (HEPN, SLATT, Primases), the way in which they operate and their evolutionary origins can be completely different. This suggests that the role of UG/Abi RTs may lie not only in the domains they contain or the genes to which they are associated but in the specific cooperation of these components in their given biological context and against specific signals that are yet to be determined.

Although there are still many other biological and mechanistic enigmas to be solved, the different UG/Abi RTs described in this work disclosed an enormous diversity of associated genes and domains. That along with the possibility of modifying non-coding RNAs and their involvement in defense functions could make the UG/Abi family RTs a prominent element in the phage-host arms race and a highly valuable source for the development of promising biotechnological and gene-editing tools.

## Acknowledgements

We are specially thankful to Rafael Pinilla-Redondo for his careful review of the manuscript and his valuable contributions and discussions.

## Funding

This work was supported by grant BIO2017-82244-P from the MCIN/ AEI/10.13039/501100011033/ FEDER “Una manera de hacer Europa” and grant PID2020-113207GB-I00 from the MCIN/ AEI /10.13039/501100011033 awarded to N.T and F.M.A. M.R.M is supported by the FPI-UAM program from Universidad Autónoma de Madrid. L.G. is supported by the Society of Fellows of Harvard University. S.A.S is a recipient of a Novo Nordisk Foundation project grant in basic bioscience [NNF18OC0052965]. A.L.B is a recipient of FPI fellowship [PRE2020-092935] from Spanish Ministry of Science and Innovation. J.I. is supported by the “Ramón y Cajal” program from the Spanish Ministry of Science [RYC-2017-22524], the Agencia Estatal de Investigación of Spain [PID2019-106618GA-I00] and the “Severo Ochoa Program for Centres of Excellence in R&D” from the Agencia Estatal de Investigación of Spain [SEV-2016-0672 (2017 to 2021) to the CBGP]. M.R.R is funded by grants from the Spanish Ministry of Science, Innovation and Universities [PGC2018-093723-A-100, AEI/FEDER] and Fundación Ramón Areces (VirHostOmics). F.Z. is supported by NIH grants [1DP1-HL141201 and 2R01HG009761-05]; the Howard Hughes Medical Institute; Open Philanthropy, the Edward Mallinckrodt, Jr. Foundation; the Poitras Center for Psychiatric Disorders Research at MIT; the Hock E. Tan and K. Lisa Yang Center for Autism Research at MIT; the Yang-Tan Molecular Therapeutics Center at McGovern, and by the Phillips family and J. and P. Poitras.

## Competing interests

Authors declare no competing interests

## Supplementary Figures

**Supplementary Figure S1.**
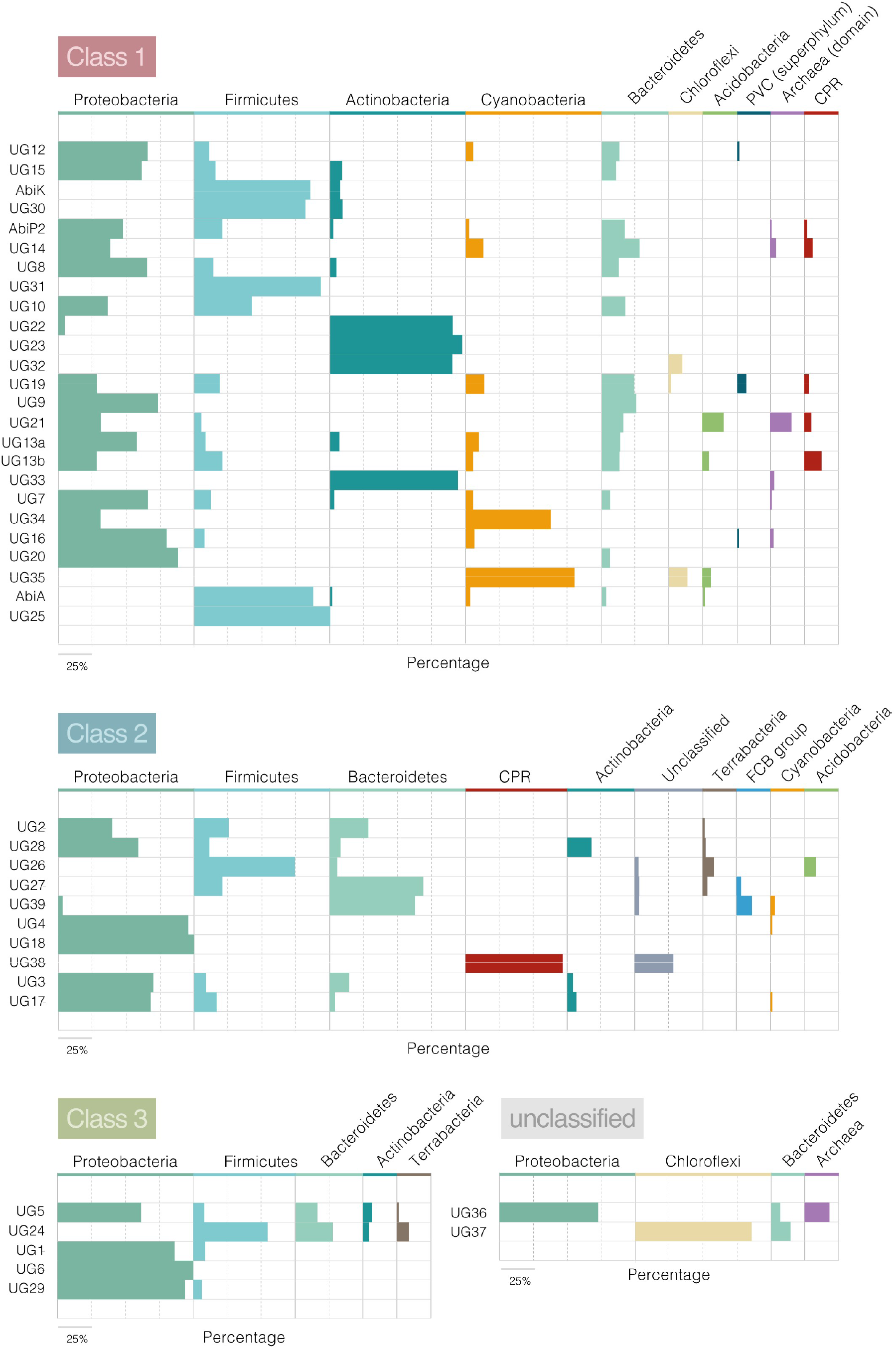
Taxonomic information of every UG/Abi RT group based on the NCBI Taxonomy database. Colors represent the different phyla in which the 5022 representative sequences are found, and bar size represents the percentage of representative sequences belonging to those phyla in each UG/Abi group. Phyla with relative abundances below 5% were removed to improve the visualization.

**Supplementary Figure S2.**
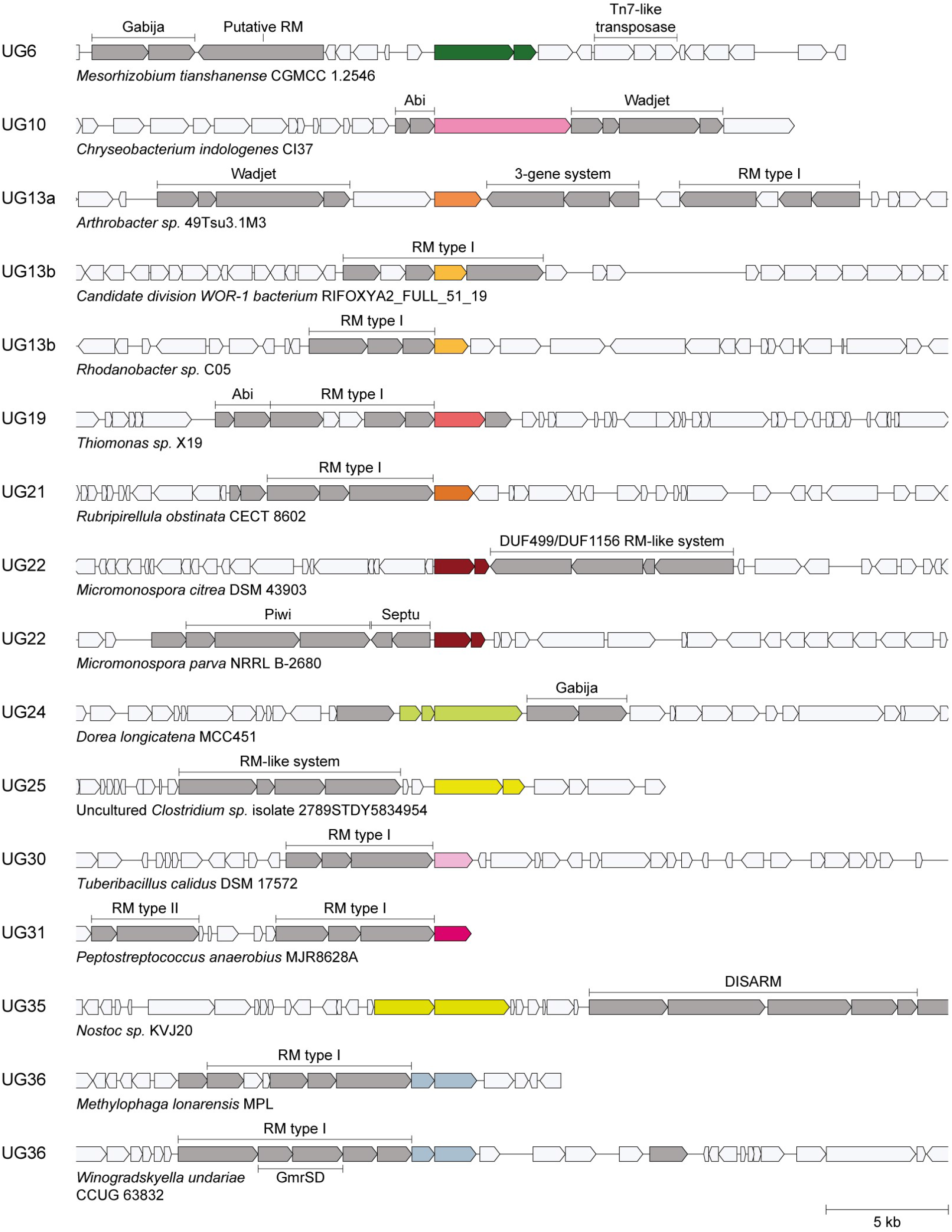
Selected genomic loci containing signatures of strong defense association next to Abi/UG RT groups with <0.15 defense association score or fewer than 50 homologs identified in the defense analysis pipeline.

**Supplementary Figure S3.**
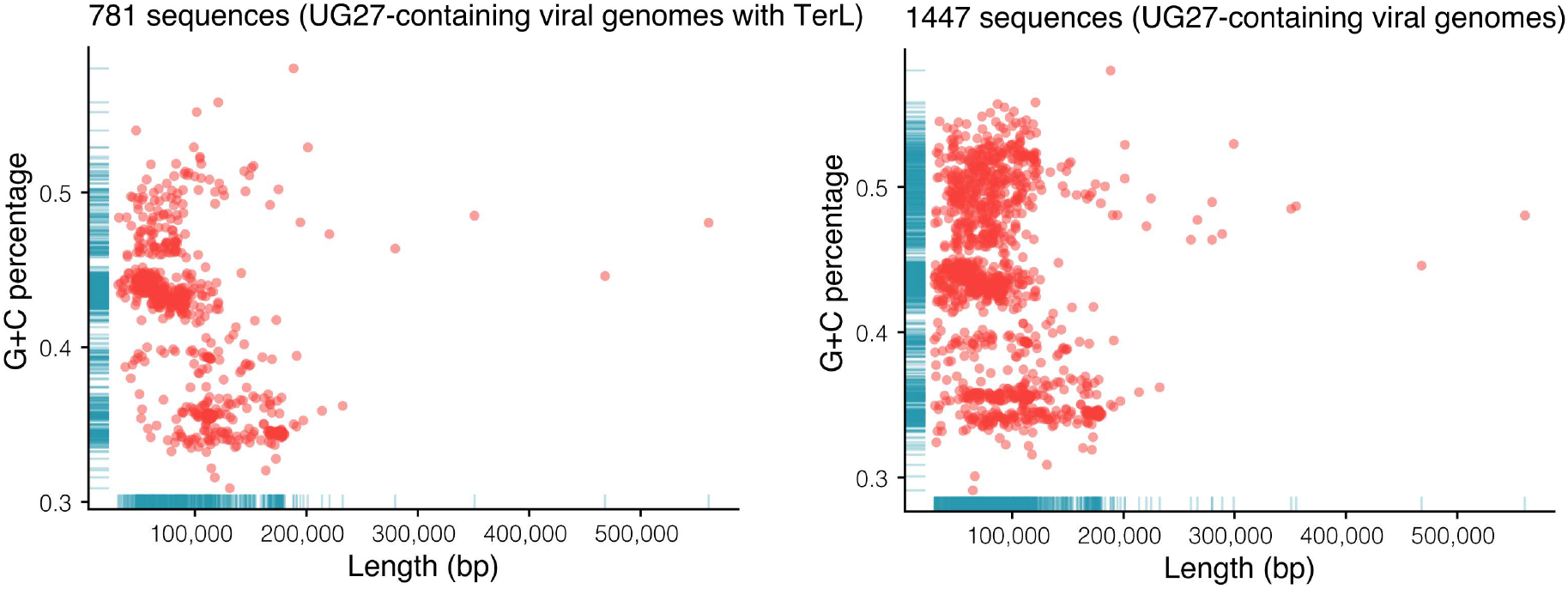
Distribution of G+C percentage and length of UG27-containing viral genomes dereplicated at 95% ANI. Left plot represents genomes obtained from IMG/VR, GPD, and mMGE databases and right plot is restricted to genomes in left plot harboring at least one TerL gene.

**Supplementary Figure S4.**
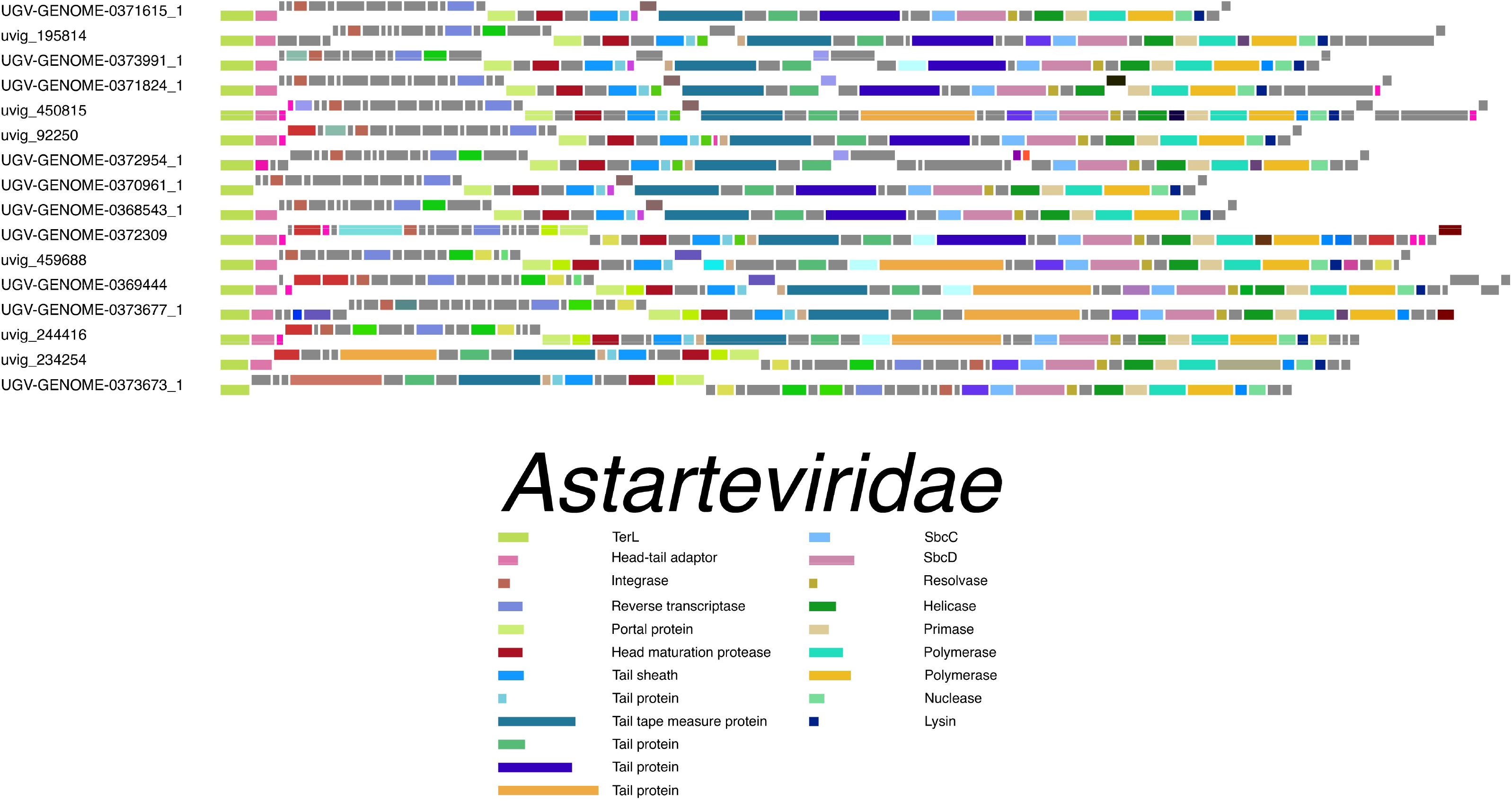
Gene maps for a representative subset of *Astarteviridae* genomes. Phage genome identifiers are given on the left. Rectangles correspond to protein-coding genes, raised or lowered based on the DNA strand they occupy. Colored genes gave matches to known phage protein domains using HHblits against the PhROGs database. The legend on the right shows the domain annotations for phage structural and replication proteins.

**Supplementary Figure S5.**
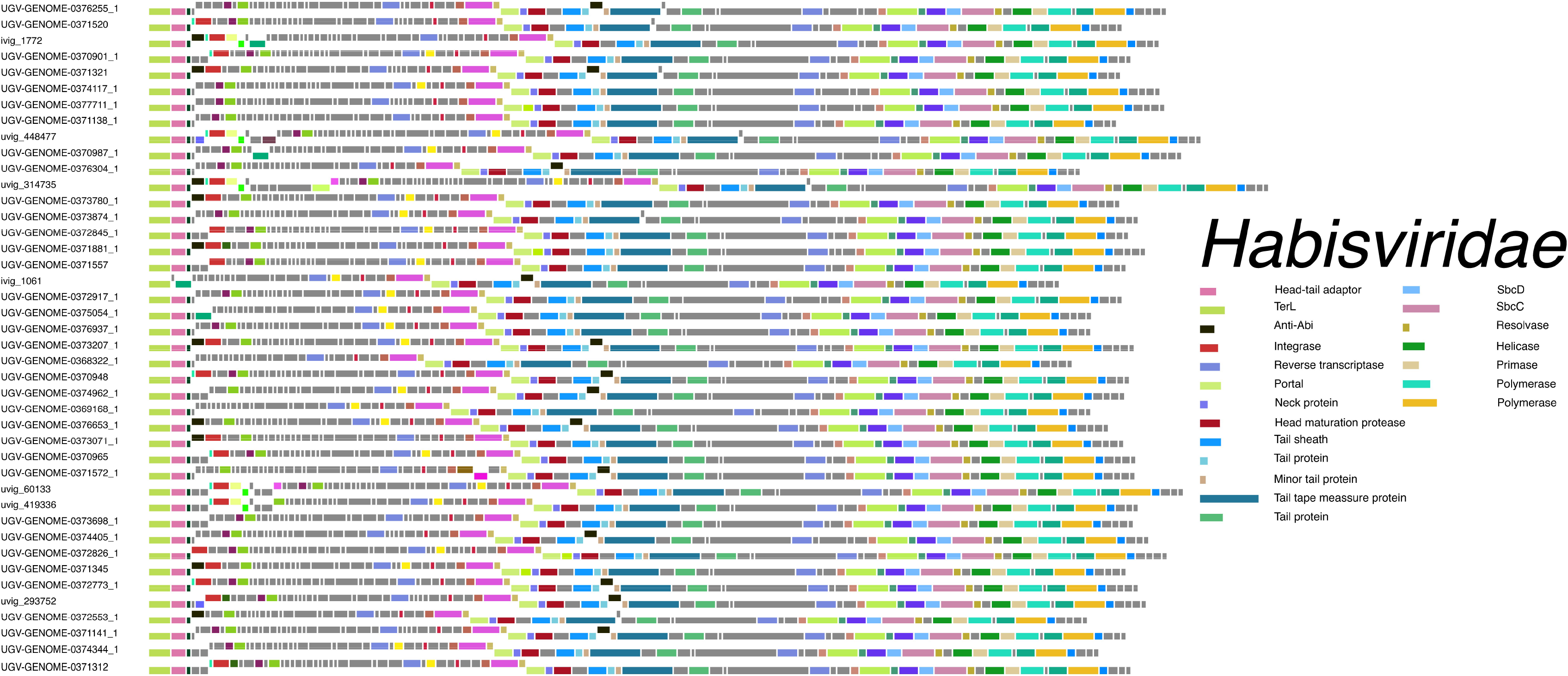
Gene maps for a representative subset of *Habisviridae* genomes. Phage genome identifiers are given on the left. Rectangles correspond to protein-coding genes, raised or lowered based on the DNA strand they occupy. Colored genes gave matches to known phage protein domains using HHblits against the PhROGs database. The legend on the right shows the domain annotations for phage structural and replication proteins.

**Supplementary Figure S6.**
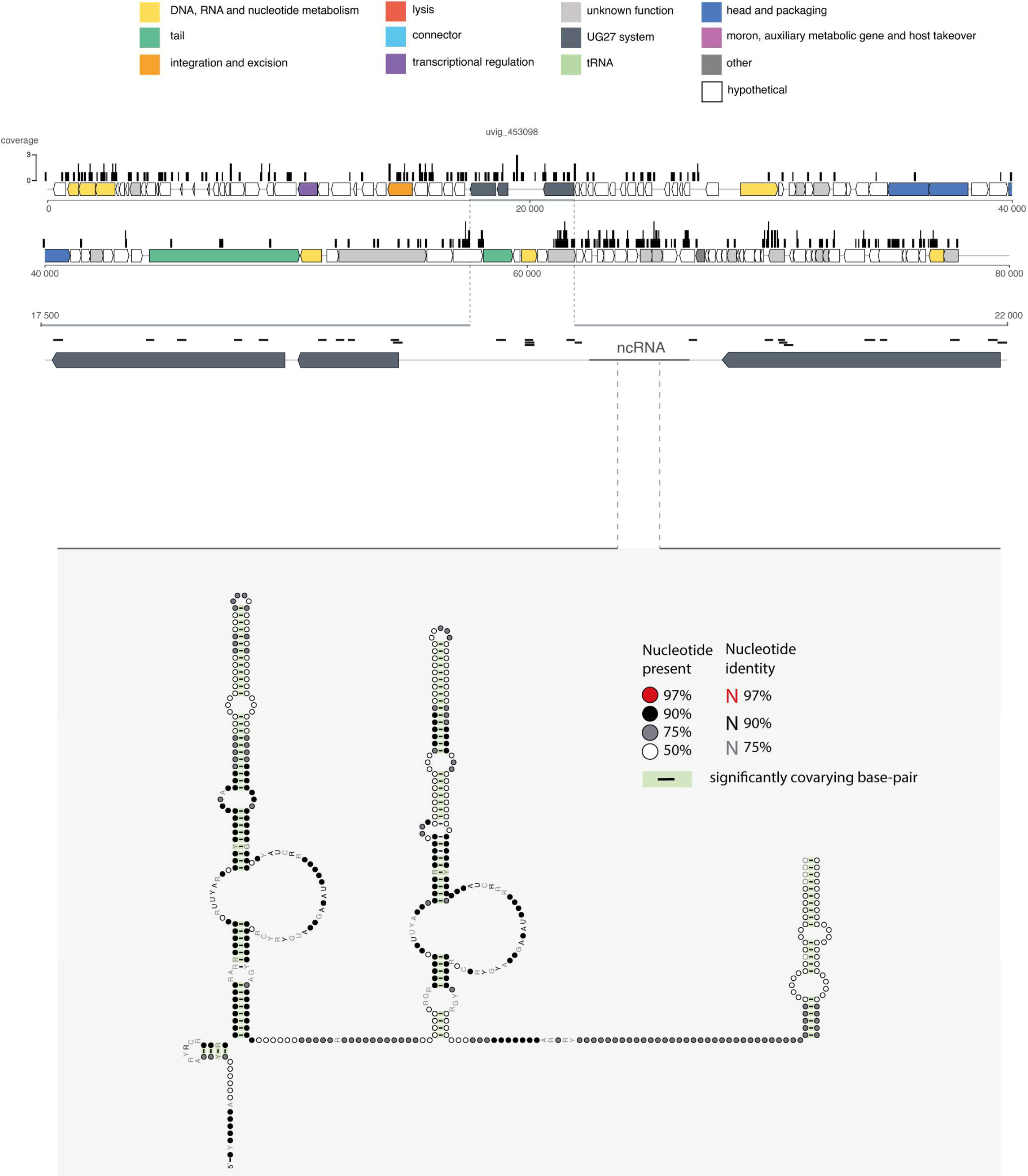
Representative UG27-containing viral genome targeted by CRISPR spacers. Annotation of the viral genome was made using PHROG profiles. Genes are colored according to PHROG functional categories. Bars above the represented genome indicate the coverage of CRISPR spacers matching the viral genome.

**Supplementary Figure S7.**
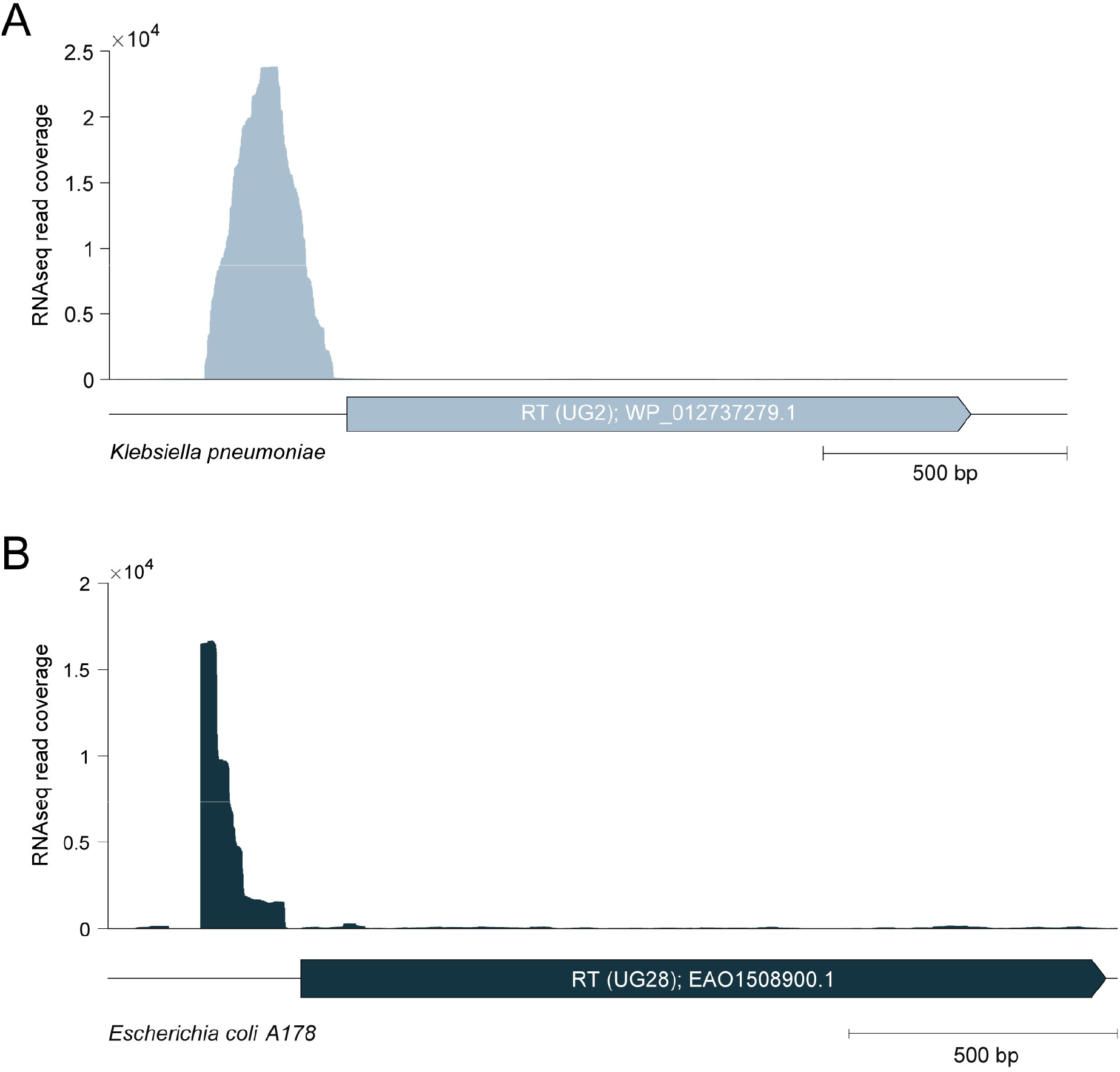
A) RNAseq read coverage for the UG2 locus from *Klebsiella pneumoniae ST23*, mapped from Sequence Read Archive SRX814863. B) RNAseq read coverage for the UG28 locus from *Escherichia coli A178*, mapped from RNA sequencing (Methods)

**Supplementary Figure S8.**
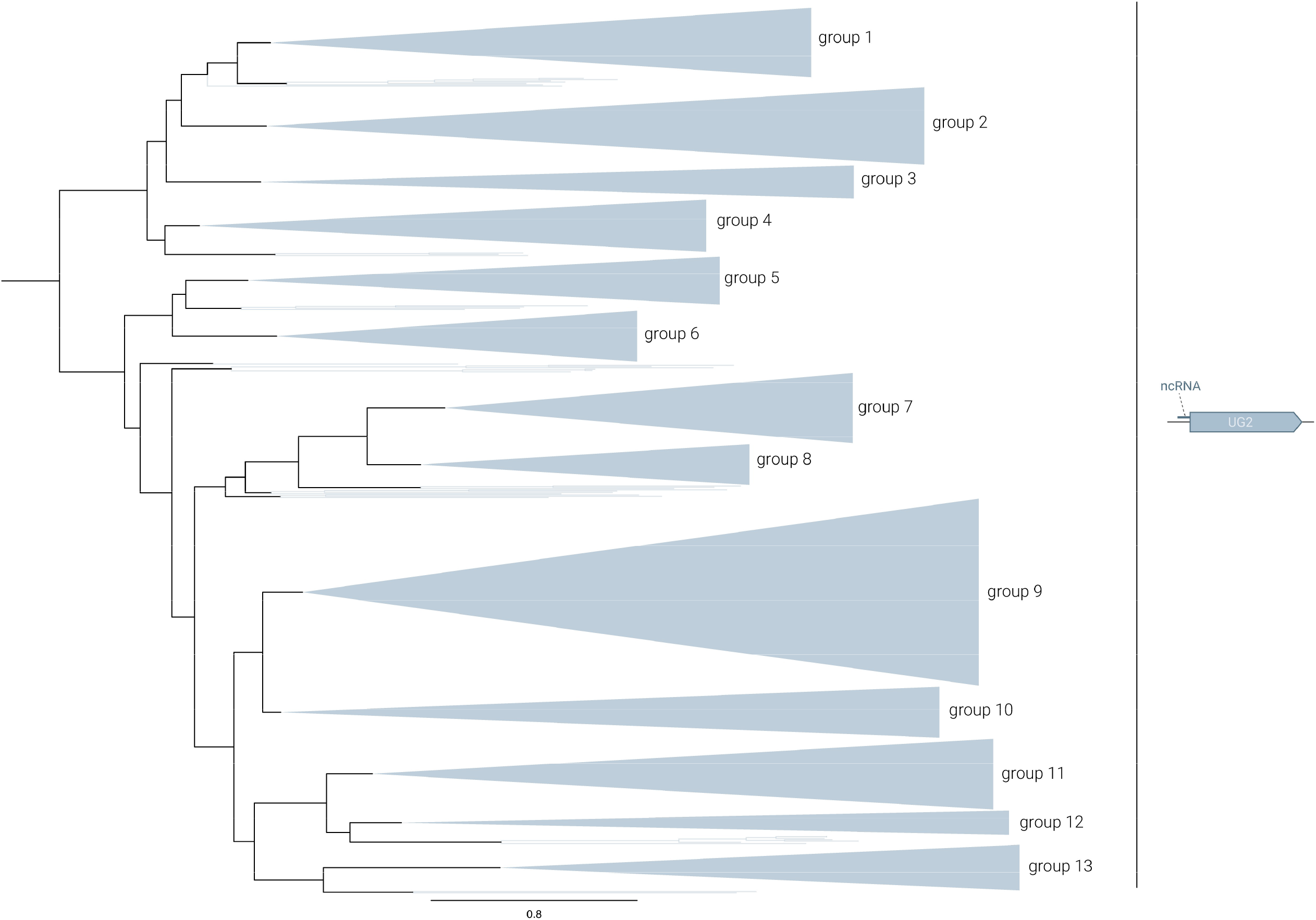
Subtree of the phylogenetic reconstruction of the 5022 representative UG/Abi RTs summarizing the different UG2 subgroups. Subgroups are collapsed based on phylogenetic and taxonomic information (Methods).

**Supplementary Figure S9.**
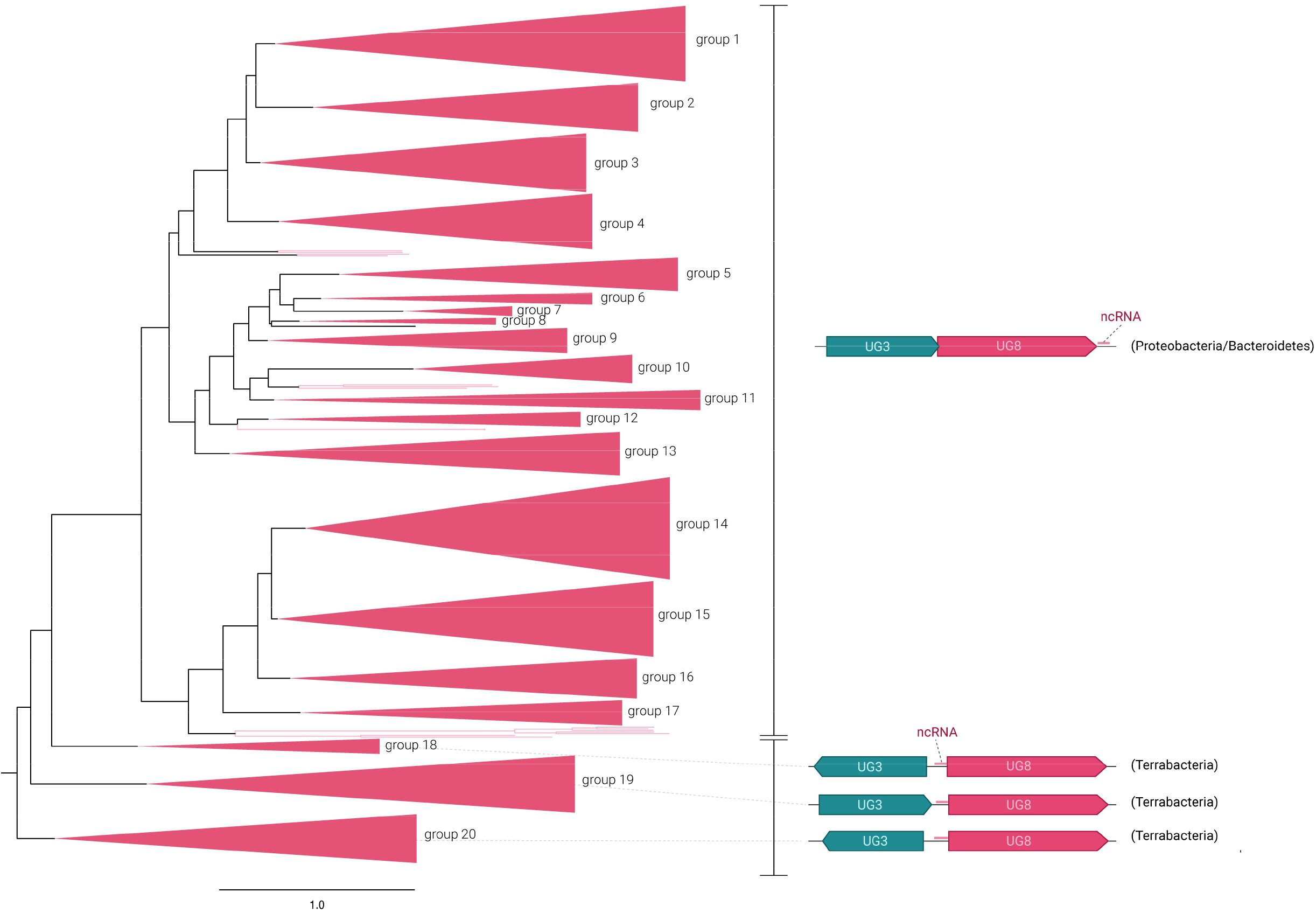
Subtree of the phylogenetic reconstruction of the 5022 representative UG/Abi RTs representing the different UG3+UG8 subgroups. Subgroups are collapsed based on phylogenetic and taxonomic information.

**Supplementary Figure S10.**
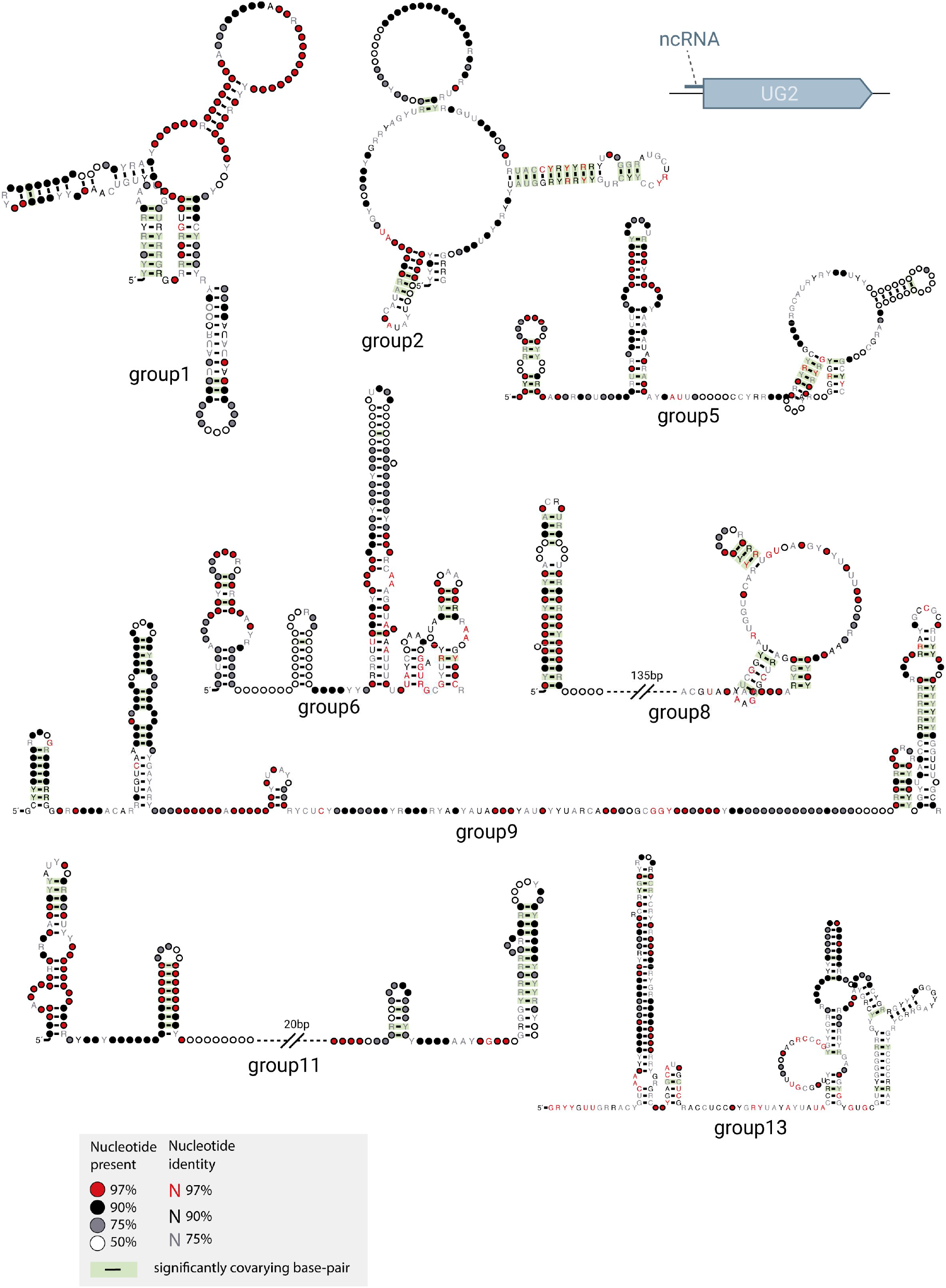
Predicted consensus structures of ncRNAs found in UG2 systems. The groups correspond to those described in Supplementary Figure S8.

**Supplementary Figure S11.**
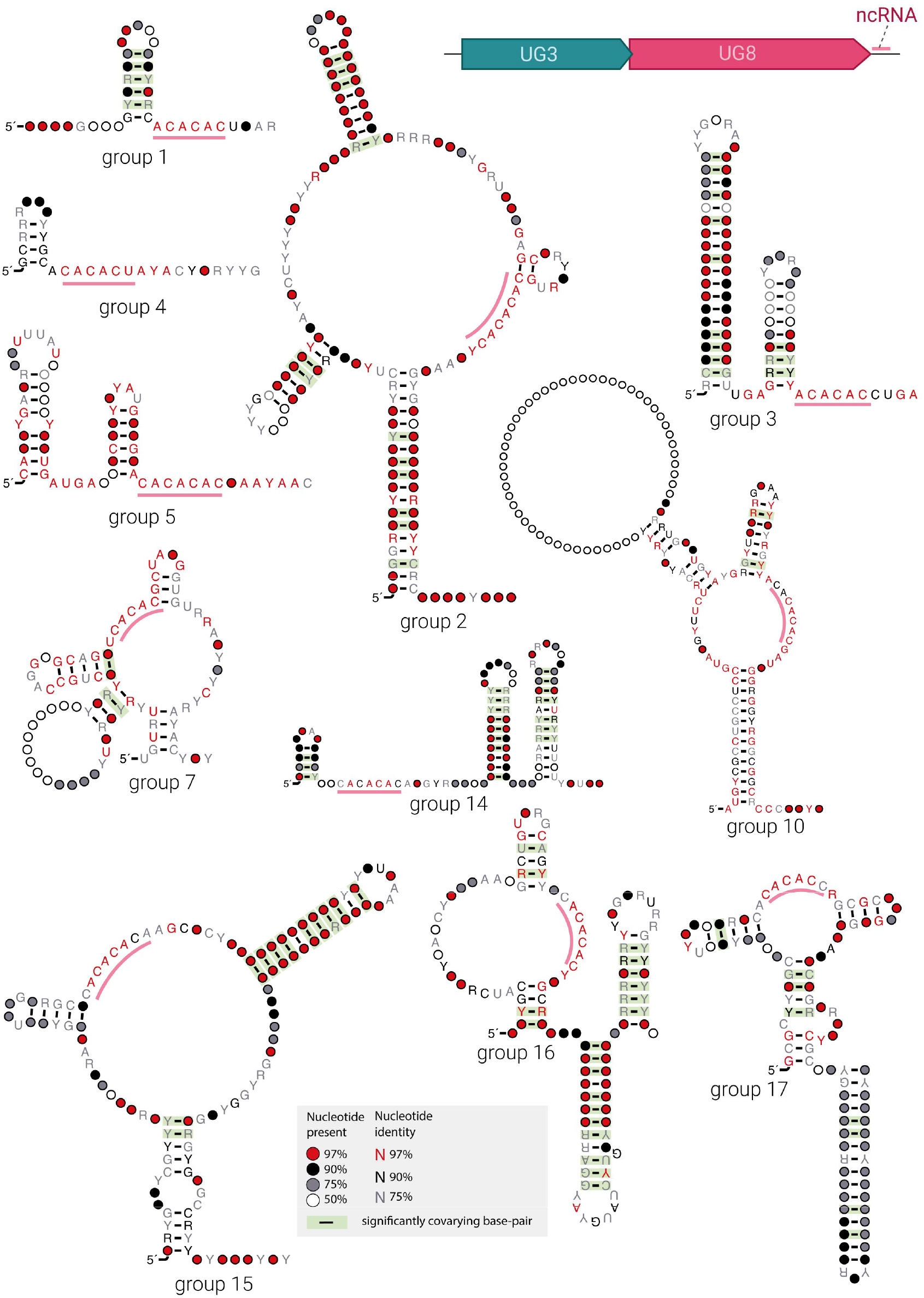
Predicted consensus structures of ncRNAs found in UG3+UG8 systems. The groups correspond to those described in Supplementary Figure S9.

**Supplementary Figure S12.**
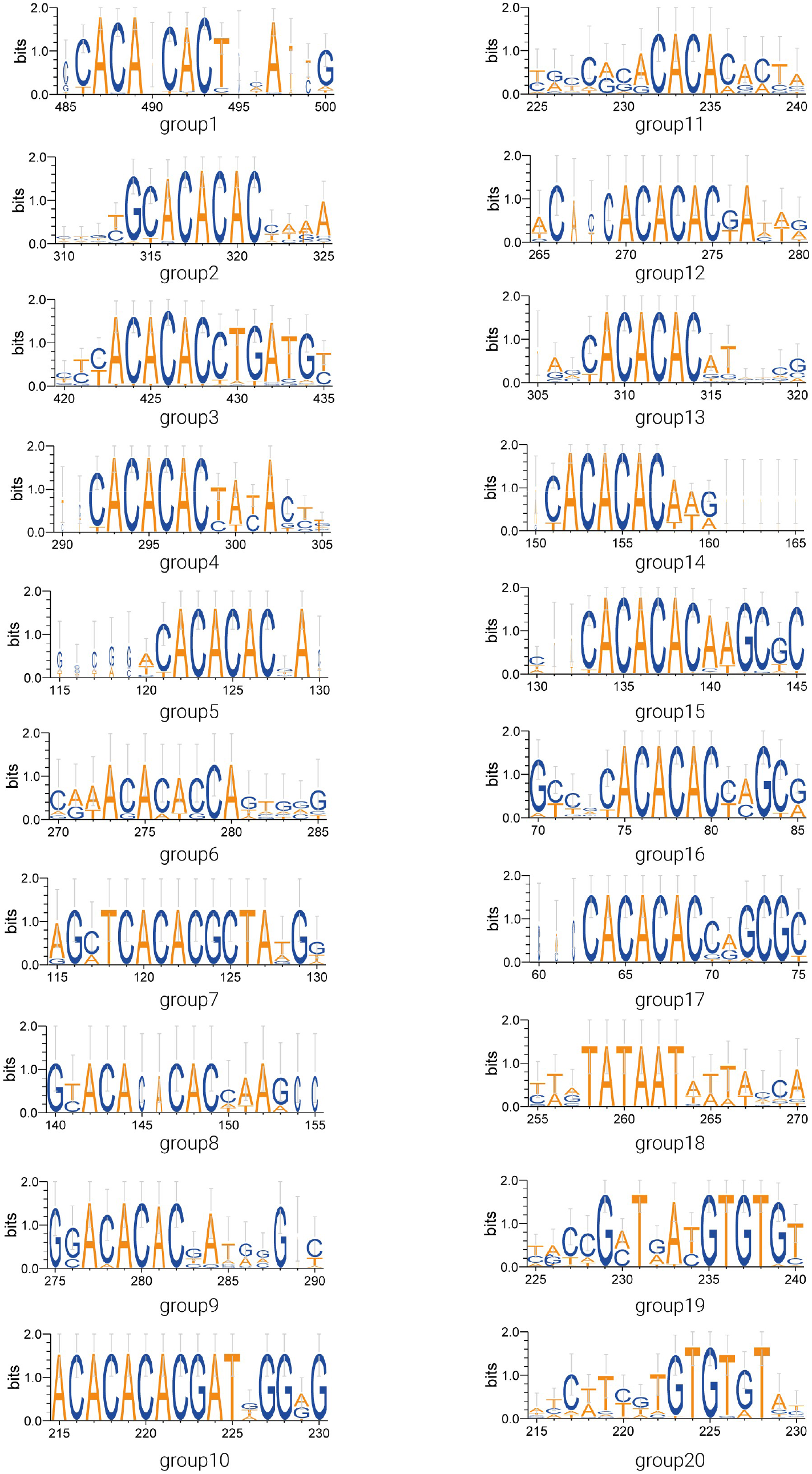
Logo representation of conserved motifs identified in predicted UG3+UG8 ncRNA sequences. The groups correspond to those described in figures S9 and S11. X-axis numbers represent aligned positions.

**Supplementary Figure S13.**
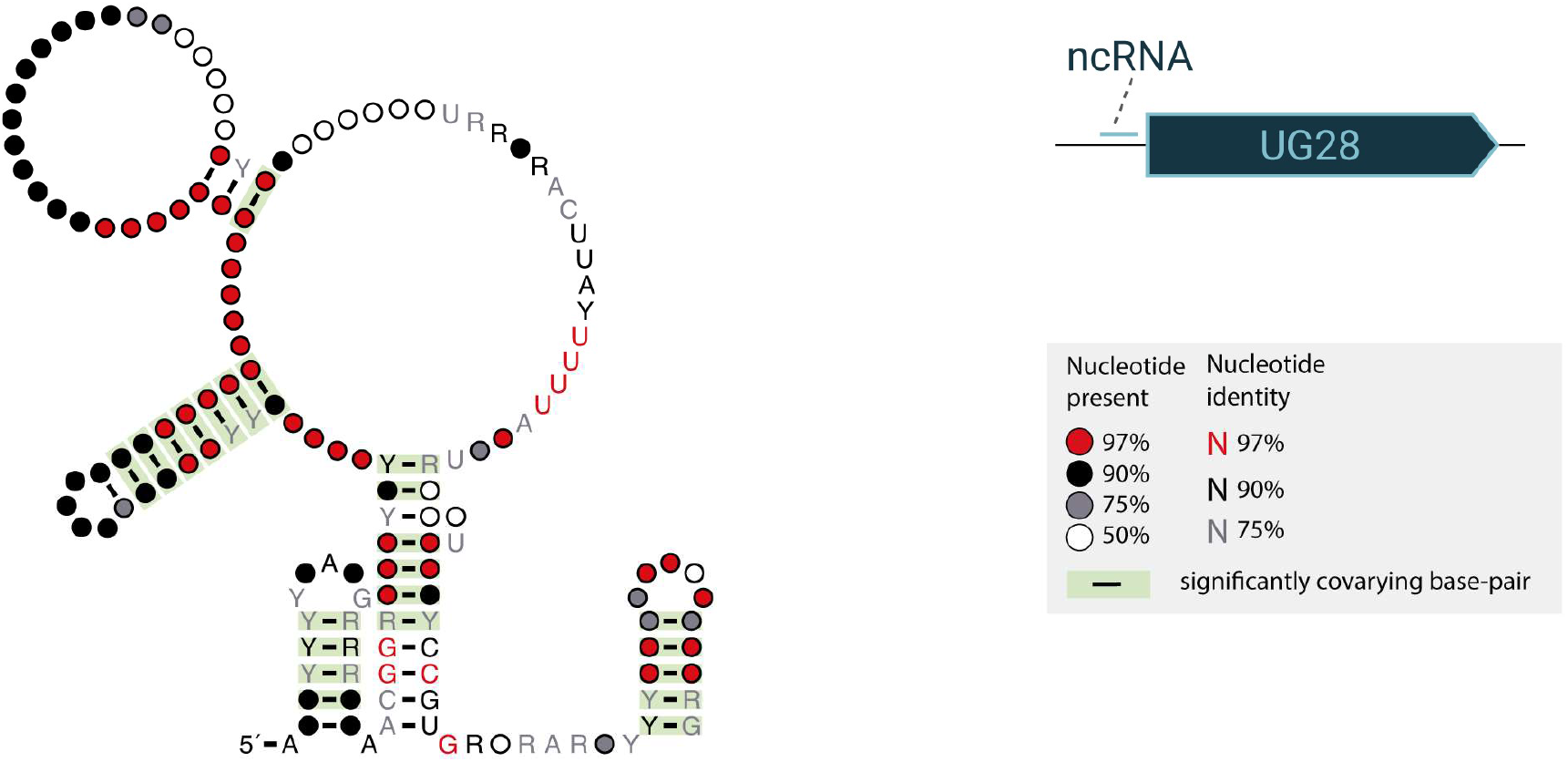
Predicted consensus structure of ncRNAs found in UG28 systems.

## Supplementary Tables and Files

**Supplementary Table S1.** Summary Table including info about representative UG/Abi RT sequences, domains, associated protein clusters, and taxonomic information.

**Supplementary Table S2.** Domain annotation of associated protein clusters.

**Supplementary Table S3.** Predicted defense scores of every UG/Abi group.

**Supplementary Table S4.** Presence of UG/Abi RTs in Mobile Genetic Elements.

**Supplementary Table S5.** Comparison of UG27-containing viral genomes with reference genomes using Average Nucleotide Identity.

**Supplementary Table S6.** Classification of phage genomes used in this study based on the APS tree

**Supplementary Table S7.** Host assignment of UG27-containing viral genomes based on CRISPR spacers matching.

**Supplementary File S1.** Phylogenetic tree in Newick format of 5022 representative UG/Abi RT sequences obtained with FastTree.

**Supplementary File S2.** Gene-sharing network of UG27-containing viral genomes and close reference sequences.

## Notes

### Competing Interest Statement

The authors have declared no competing interest.

